# Comparative GWAS identifies a role for Mendel’s green pea gene in the nonphotochemical quenching kinetics of sorghum, maize, and arabidopsis

**DOI:** 10.1101/2023.08.29.555201

**Authors:** Seema Sahay, Nikee Shrestha, Henrique Moura Dias, Ravi V. Mural, Marcin Grzybowski, James C. Schnable, Katarzyna Głowacka

## Abstract

Photosynthetic organisms must cope with rapid fluctuations in light intensity. Nonphotochemical quenching (NPQ) enables the dissipation of excess light energy as heat under high light conditions, whereas its relaxation under low light maximizes photosynthetic productivity. We quantified variation in NPQ kinetics across a large sorghum (*Sorghum bicolor*) association panel in four environments, uncovering significant genetic control for NPQ. A genome-wide association study (GWAS) identified 20 unique regions in the sorghum genome associated with NPQ. We detected strong signals from the sorghum ortholog of *Arabidopsis thaliana SUPPRESSOR OF VARIEGATION3* (*SVR3*) involved in plastid–nucleus signaling and tolerance to cold. By integrating GWAS results for NPQ across maize (*Zea mays*) and sorghum association panels, we identified a second gene, *NON-YELLOWING 1* (*NYE1*), originally identified by Gregor Mendel in pea (*Pisum sativum*) and involved in the degradation of photosynthetic pigments in light-harvesting complexes, along with *OUTER ENVELOPE PROTEIN 37* (*OEP37*), that encodes a transporter in chloroplast envelope. Analysis of *nye1* insertion alleles in *A. thaliana* confirmed the effect of this gene on NPQ kinetics across monocots and eudicots. We extended our comparative genomics GWAS framework across the entire maize and sorghum genomes, identifying four additional loci involved in NPQ kinetics. These results provide a baseline for engineering crops with improved NPQ kinetics and increasing the accuracy and speed of candidate gene identification for GWAS in species with high linkage disequilibrium.

## Introduction

The leaves of plants grown under field conditions experience rapid fluctuations in the quantity of photosynthetically active light reaching their surface, resulting from both intermittent clouds passing in front of the sun and moving shadows caused by the wind blowing through leaf canopies (Tang *et al*. 1988; Burgess *et al*. 2016). In the absence of buffering mechanisms, rapid increases in light intensity can produce singlet oxygen and other reactive oxygen species (ROS) that can damage both the photosystems and other proteins throughout the cell (Muller *et al*. 2001). Plants have evolved several mechanisms to safely dissipate excess energy absorbed by the photosynthetic apparatus as heat, minimizing the formation of damaging ROS due to changes in light intensity. These mechanisms are collectively known as nonphotochemical quenching (NPQ). In discussions of NPQ below, we focus specifically on two components of NPQ: energy-dependent quenching (qE) and zeaxanthin-dependent quenching (qZ). qE reacts within one-tenth of a second (Muller *et al*. 2001)—and is, therefore, the mechanism that responds most rapidly to sudden increases in light intensity—whereas qZ has a slower relaxation time of 10 to 15 minutes. Several proteins involved in NPQ have been identified through forward genetic screens, including *PHOTOSYSTEM II SUBUNIT S* (*PSBS*, also named *NPQ4* based on the corresponding mutant; Li *et al*. (2000)), *VIOLAXANTHIN DE-EPOXIDASE 1* (*VDE1*; or *NPQ1*), and *ZEAXANTHIN EPOXIDASE* (*ZEP*; or *NPQ2*) (Niyogi *et al*. 1998).

The kinetics of NPQ induction and relaxation reflect a core fitness trade-off for plants. Failure to induce NPQ rapidly enough or the inability to reach sufficient maximum values under high light conditions exposes plants to the risk of substantial damage and diminished photosynthetic productivity due to photoinhibition. However, NPQ induction beyond the levels necessary to protect plants from damage leads to the wasteful dissipation of photosynthetic energy as heat and reduced productivity, as does an insufficiently rapid drop in NPQ when light intensity declines. Optimal and maximal values for NPQ induction and relaxation likely vary depending on both the external environment (e.g., frequency and type of cloud cover or maximum light intensity during the growing season) and properties of plant canopy architecture (Tang *et al*. 1988; Pearcy 1990; Burgess *et al*. 2016; Kaiser *et al*. 2018). Whereas natural selection has shaped the response of plants to fluctuating light intensities over hundreds of millions of years, key crops have been under selection for performance in the artificial environments of agricultural fields for only thousands of years and have experienced selection for performance at modern planting densities that have profoundly reshaped crop and canopy architectures for only decades. As a result, the kinetics of NPQ in major crops may not be optimized for current growing environments (Zhu *et al*. 2004). Transgenic interventions to alter NPQ kinetics have achieved increases in dry biomass accumulation in tobacco (*Nicotiana tabacum*) plants grown under field conditions of 14-20% (Kromdijk *et al*. 2016) and significant increases in soybean (*Glycine max*) yields in one of two environments tested (De Souza *et al*. 2022).

Both crops and wild species exhibit substantial natural variation in NPQ (Jung and Niyogi 2009; Rungrat *et al*. 2019; Gamba *et al*. 2022; Wei *et al*. 2022). A study of the maximum NPQ induction in rice (*Oryza sativa*) identified 33 loci, including the rice ortholog of *PSBS* (Wang *et al*. 2017). In soybean, a study of variation in the epoxidation state of the xanthophyll pigments, a key component of NPQ identified 15 loci, including one proximal to the soybean ortholog of *VDE* (Herritt *et al*. 2016). A genome-wide association study (GWAS) conducted in *Arabidopsis thaliana* identified 15 loci associated with variation in NPQ induction, maximum value, or relaxation, including one which co-localized with the previously characterized *PSBS* (Rungrat *et al*. 2019). *A. thaliana*, rice, and soybean are all C_3_ plants. A recent GWAS in maize (*Zea mays*), a C_4_ plant identified 18 confident associated regions with different traits describing the kinetics of NPQ induction, maximum value, or relaxation, including a signal associated with the maize ortholog of *PSBS*. In six cases, insertion mutants in the *A. thaliana* orthologs of candidate genes adjacent to maize GWAS hits produced similar alterations in NPQ kinetics to those observed in maize (Sahay *et al*. 2023).

Here, we sought to identify genetic loci associated with variation in NPQ kinetics across a widely characterized and recently re-sequenced sorghum association panel (*Sorghum bicolor*) (Casa *et al*. 2008; Mural *et al*. 2021; Boatwright *et al*. 2022), a drought-tolerant and nitrogen-use-efficient relative of maize that is widely grown as a staple crop in Subsaharan Africa and South Asia. We identified several significant NPQ trait-associated loci including the sorghum ortholog of a gene previously reported in maize and *A. thaliana* (Sahay *et al*. 2023) and signal from sorghum and maize orthologs of *NON-YELLOWING 1 (NYE1)*. Multiple insertion alleles of *nye1* exhibit impaired NPQ, consistent with the maize and sorghum phenotypes. Collectively, these results begin to address the historically limited characterization of natural variation NPQ among C_4_ species and support a core set of existing functional proteins playing conserved roles in determining NPQ kinetics across angiosperms.

## Results

We observed substantial variation in nine traits describing NPQ kinetics (Table 1) in a population of 339 sorghum genotypes grown under low-nitrogen (LN) conditions in 2020 (Figure 1, Supplemental Figure S1). While the LN treatment was sufficient to decrease sorghum grain yield by 48% relative to high-nitrogen (HN) controls, and other traits measured in the same field experiment, such as flag leaf width and length, also showed significant differences between treatments (Grzybowski *et al*. 2022), we did not observe consistent differences in NPQ kinetics between LN and HN conditions years (Supplemental Figure S2). NPQ kinetics were more correlated between different nitrogen conditions in the same year than between the same nitrogen conditions in different years, consistent with environmental factors other than access to nitrogen playing a greater role in shaping variation in NPQ (Supplemental Figure S3). The average within-environment heritability of NPQ kinetic traits was modestly lower in this sorghum experiment than previously reported for maize traits (Table 2). Given the lack of clear response to nitrogen supply, we focused on measurements of NPQ kinetics from the LN conditions from 2020, as traits exhibited the higher average heritability in this environment (H^2^ = 0.50) than the other three from which data was collected (H^2^ = 0.42, 0.40 and 0.35, Table 2).

**Figure 1.**
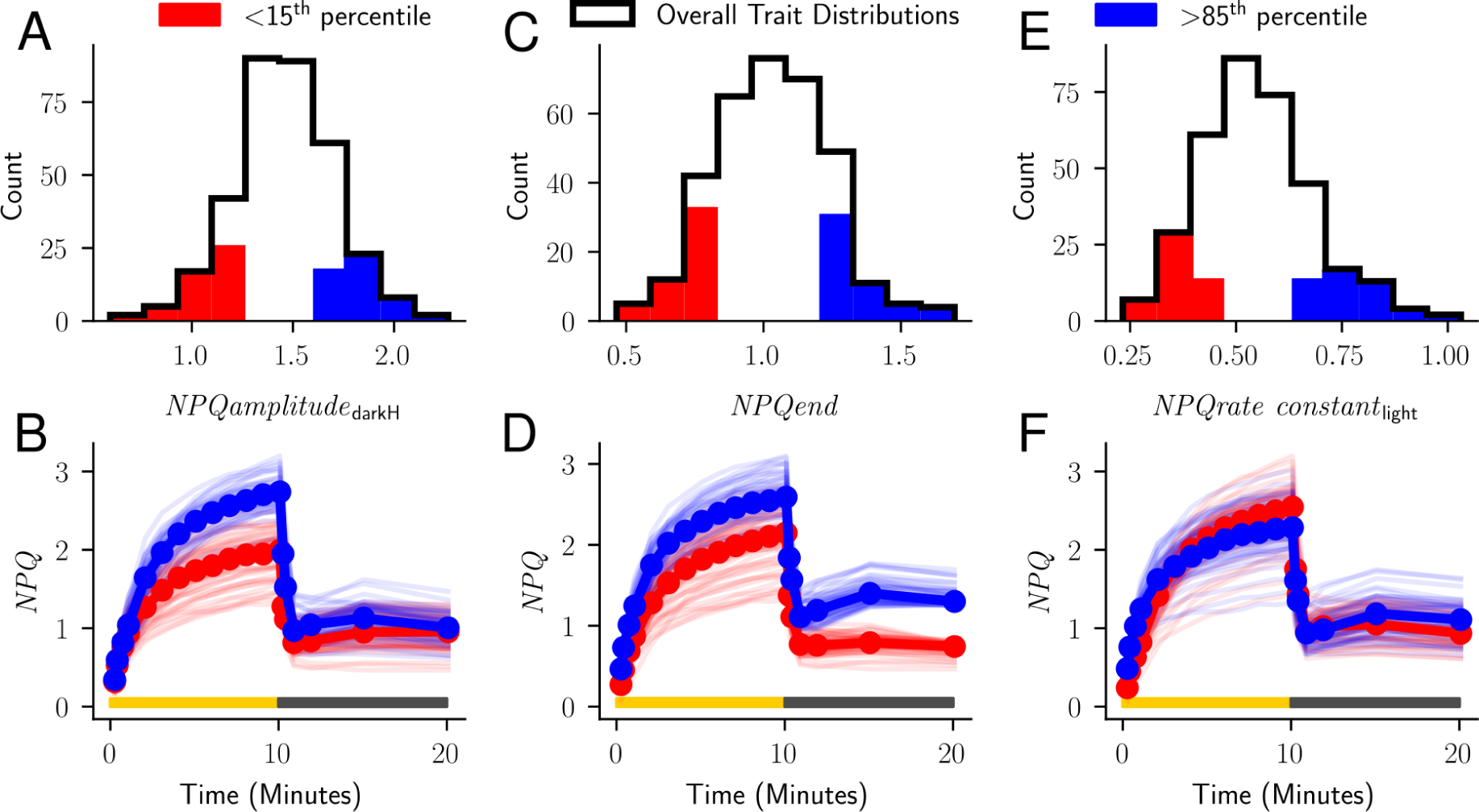
Patterns of NPQ kinetics associated with phenotypic extremes for three selected NPQ kinetics traits. **A)** Distribution of observed values for *NPQamplitude_darkH_* among the genotypes of the sorghum association panel grown in Lincoln, Nebraska, USA in the summer of 2020 under low-nitrogen (LN) conditions (n=339 sorghum geno-types). **B)** Genotype-specific patterns of NPQ induction and relaxation for the 15% of sorghum genotypes with the lowest *NPQamplitude_darkH_* scores (light red lines) and the 15% of sorghum genotypes with the highest *NPQamplitude_darkH_* scores (light blue lines). Median NPQ response among sorghum genotypes with the lowest and highest *NPQamplitude_darkH_* shown in solid red and blue, respectively. **C)** Distribution of observed values for *NPQ_end_* among the same set of sorghum lines shown in A (n=339 sorghum genotypes). **D)** Genotype-specific and median NPQ induction and relaxation curves for sorghum genotypes at both extremes of the distribution for *NPQ_end_*, presented as described in B. **E)** Distribution of observed values for *NPQrate constant_light_* among the same set of sorghum lines shown in A. One genotype with median *NPQrate constant_light_* >1.5 was excluded to improve figure readability (n=338 sorghum genotypes). **F)** Genotype-specific and median NPQ induction and relaxation curves for sorghum genotypes at both extremes of the distribution for *NPQrate constant_light_*, presented as described in B. Data used to produce this figure is given in Supplemental Data Sets S1A.

**Table 1.** Descriptions of nonphotochemical quenching (NPQ) kinetic traits investigated in this paper.

| Trait Name | Trait Description |
| --- | --- |
| $NPQ_{slope_{lightH}}$ | How fast NPQ is induced under high light. Estimated from a curve fit to the different measurements of NPQ collected during the 10-minutes high-light treatment. |
| $NPQ_{rate\ constant_{light}}$ | Estimate of the rate at which NPQ reaches 63% of its final maximum potential value after being exposed to high light. |
| $NPQ_{asymptote_{lightH}}$ | The maximum potential value NPQ would reach under prolonged high light. Estimated from a curve fit to the different measurements of NPQ collected during the 10-minutes high-light treatment. |
| $NPQ_{start_{darkH}}$ | The NPQ value at the transition point between high light and dark treatments. Estimated from a curve fit to the different measurements of NPQ collected during the 10-minutes dark treatment. |
| $NPQ_{slope_{darkH}}$ | How fast NPQ relaxes once the lights are turned off. Estimated from a curve fit to the different measurements of NPQ collected during the 10-minute dark treatment. |
| $NPQ_{rate\ constant_{dark}}$ | Estimate of the rate at which NPQ reaches 63% of its final minimum potential value after the end of illumination. |
| $NPQ_{amplitude_{darkH}}$ | Size of the change between estimated maximum NPQ induction and maximum post-induction NPQ relaxation. |
| $NPQ_{residual_{dark}}$ | The amount of potential NPQ relaxation beyond that which occurs during the 10 minute dark treatment. |
| $NPQ_{end}$ | Remaining NPQ still present after 10 minutes in the dark. |

**Table 2.** Heritability of non-photochemical quenching (NPQ) traits in sorghum and maize (LN: Low nitrogen condition and HN: High nitrogen condition).

| Trait | Sorghum H<sup>2</sup> |  |  |  | Maize H<sup>2</sup> |  |
| --- | --- | --- | --- | --- | --- | --- |
|  | LN2020 | LN2021 | HN2020 | HN2021 | 2020 | 2021 |
| *NPQslope<sub>lightH</sub>* | 0.43 | 0.37 | 0.39 | 0.33 | 0.60 | 0.59 |
| *NPQrate constant<sub>light</sub>* | 0.51 | 0.34 | 0.44 | 0.16 | 0.60 | 0.60 |
| *NPQasymptote<sub>lightH</sub>* | 0.54 | 0.40 | 0.44 | 0.48 | 0.56 | 0.50 |
| *NPQstart<sub>darkH</sub>* | 0.46 | 0.27 | 0.39 | 0.46 | 0.55 | 0.51 |
| *NPQslope<sub>darkH</sub>* | 0.55 | 0.49 | 0.30 | 0.41 | 0.62 | 0.61 |
| *NPQrate constant<sub>dark</sub>* | 0.51 | 0.47 | 0.37 | 0.34 | 0.63 | 0.54 |
| *NPQamplitude<sub>darkH</sub>* | 0.44 | 0.29 | 0.42 | 0.40 | 0.56 | 0.50 |
| *NPQresidual<sub>dark</sub>* | 0.54 | 0.26 | 0.51 | 0.51 | 0.49 | 0.45 |
| *NPQ<sub>end</sub>* | 0.53 | 0.24 | 0.50 | 0.49 | 0.53 | 0.48 |
| **Average** | **0.57** | **0.53** | **0.50** | **0.35** | **0.42** | **0.40** |

A set of 25 significant marker-trait associations for five NPQ-related traits were identified at a threshold of *≥* 0.1 resampling model inclusion probability (RMIP) in a FarmCPU GWAS with resampling analysis conducted with 4.4 million resequencing derived markers (Figure 2A). The 25 marker-trait associations identified formed 20 unique clusters because in some cases the same or adjacent genetic markers in high linkage disequilibrium (LD) were identified across multiple NPQ kinetic traits. We identified fewer significant associations in the analysis of NPQ kinetics traits scored in the three environments with lower heritability (Supplemental Figure S4).

**Figure 2.**
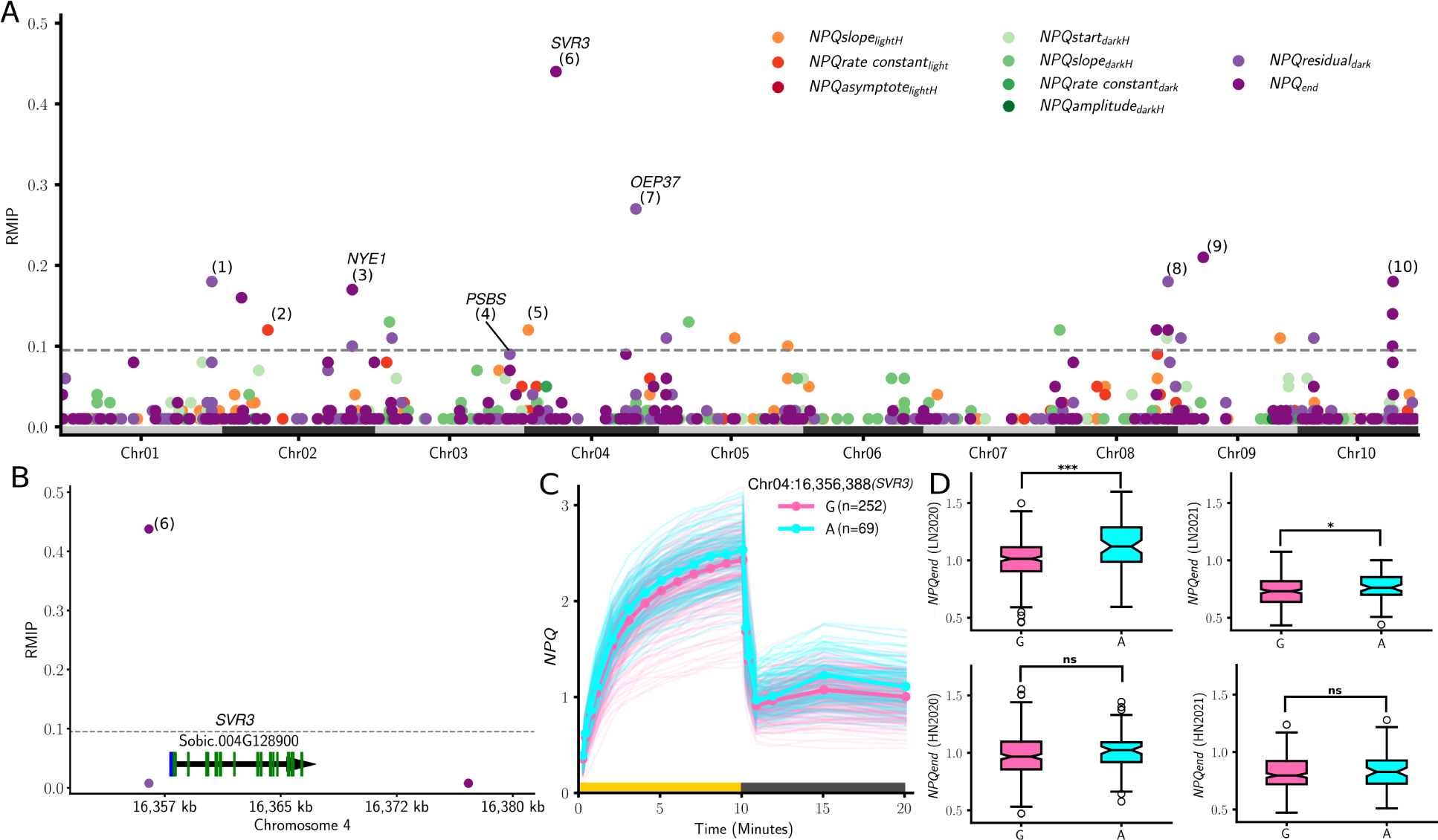
Results of genome-wide associations for NPQ kinetic traits in sorghum. **A)** Resampling model inclusion probabilities (RMIPs) for genetic markers linked to variation in nine sorghum NPQ traits (Table 1) measured in the sorghum association panel grown under low-nitrogen conditions in the summer of 2020. The dashed horizontal gray line separates points meeting the RMIP threshold of *≥* 0.10 from those with RMIP *≤* 0.09. Numbers in parenthesis indicate the order of trait-associated SNPs discussed in Supplemental Table S1. The patterns of change in NPQ response curves associated with each of these nine traits are shown in Supplemental Figure S1. **B)** Zoomed in view of trait-associated SNPs from panel A for a 30-kb region including the highest RMIP marker located at 16,356,388 on sorghum chromosome 4. Green boxes indicate exons. Blue boxes indicate untranslated regions of exons. **C)** NPQ response curves for sorghum genotypes homozygous for either the major allele (G) or the minor allele (A) Chr04:16,356,388. **(D)** *NPQ_end_* values measured under low-nitrogen conditions (LN2020 and LN2021) and high-nitrogen conditions (HN2020 and HN2021) for sorghum geno-types homozygous for either G or A at Chr04:16,356,388. In panel D colored boxes indicate the range from the 25th-75th percentile of values, black lines with notches within the colored boxes indicate the median value, whiskers indicate the most extreme values within 1.5x the interquartile range and white circles with black outlines indicate the values of datapoints outside that range. For each comparison in D, n=252 sorghum genotypes for the G allele and n=69 sorghum genotypes for the A allele. * = *P* <0.05, ** = *P* <0.001, n.s. = not significant (two-tailed t-test). Data used to produce this figure are given in Supplemental Data Sets S1, S2, and S3A.

The most consistently identified signal in the GWAS came from a marker located at position 16,356,388 on sorghum chromosome 4. This marker is 1.4 kb upstream from Sobic.004G128900 (Figure 2B) more than 48 kb from the next closest gene (Supplemental Figure S5). Sorghum genotypes homozygous for the minor allele at this locus exhibited more rapid induction of NPQ and higher maximum NPQ kinetics after 10 minutes under high-light conditions, but slower relaxation of NPQ under dark conditions than genotypes homozygous for the major allele (Figure 2C). Sobic.004G128900 is the sole sorghum ortholog of the single-copy *A. thaliana* gene *SUPPRESSOR OF VARIEGATION3* (*SVR3*, At5g13650) encoding a protein involved in ROS signaling and tolerance to light intensity stress (Liu *et al*. 2010; Saini *et al*. 2011). *A. thaliana* loss-of-function mutants of *SVR3* accumulate significantly less of the photosystem II (PSII) reaction center protein D1, suggesting a potential mechanism by which alteration in the function of this protein might affect NPQ kinetics (Liu *et al*. 2010). We also observed significant differences between the *NPQ_end_* for genotypes homozygous for the two possible alleles at marker position 16,356,388 on sorghum chromosome 4 under LN conditions in both 2020 and 2021 (*p* <0.001 and *p* <0.03), but not in the HN treatment in either year (Figure 2D).

The second most consistently identified signal was located at position 56,606,003 on sorghum chromosome 4, approximately 500 kb from the sorghum ortholog of Zm00001d017171 (Sobic.004G210500), a maize gene identified in a GWAS of NPQ kinetic traits. Both Sobic.004G210500 and Zm00001d017171 are orthologous to *OUTER ENVELOPE PROTEIN 37* (*OEP37*, At2g43950), an *A. thaliana* gene encoding a chloroplast-localized ion membrane channel protein, a null allele of which shows altered NPQ kinetics (Sahay *et al*. 2023). Sobic.004G210500 was modestly outside the mapping interval defined by linkage disequilibrium for this GWAS hit, 555 kb away from the identified GWAS hit (Supplemental Table S1). While functional variants in *PSBS* have been linked to variation in maximum NPQ across rice, *A. thaliana*, and maize (Wang *et al*. 2017; Rungrat *et al*. 2019; Sahay *et al*. 2023), the closest signal to the sorghum *PSBS* gene (Sobic.003G370000) was slightly more than 1 Mb away and modestly below our RMIP threshold for significance (Supplemental Table S1).

Two significantly trait-associated genetic markers on sorghum chromosome 2, separated by 21 kb were associated with traits describing relaxation of NPQ in the dark following challenge with high light: Chr02:65,879,368 (*NPQresidual_dark_*, RMIP = 0.10) and Chr02:65,900,630 (*NPQ_end_*, RMIP = 0.17). These markers are located in a region of elevated LD, with the window of genetic markers exhibiting LD *≥* 0.25 with Chr02:65,900,630 encompassing the interval 65,501,078–66,013,031 bp (511 kb), containing 67 annotated gene models (Figure 3A). We considered 45 of these as unlikely candidates based on a lack of expression in sorghum leaf tissue; we excluded another 11 gene models with evidence of expression in leaves, but lacked functional annotations or had functional annotations inconsistent with a role in the kinetics of NPQ. However, two genes in the mapping interval had plausible functional links to photosynthesis. Another eight genes had functional annotations that, while not strongly linked to photosynthesis or NPQ, were also consistent with a role in NPQ or could not be ruled out.

**Figure 3.**
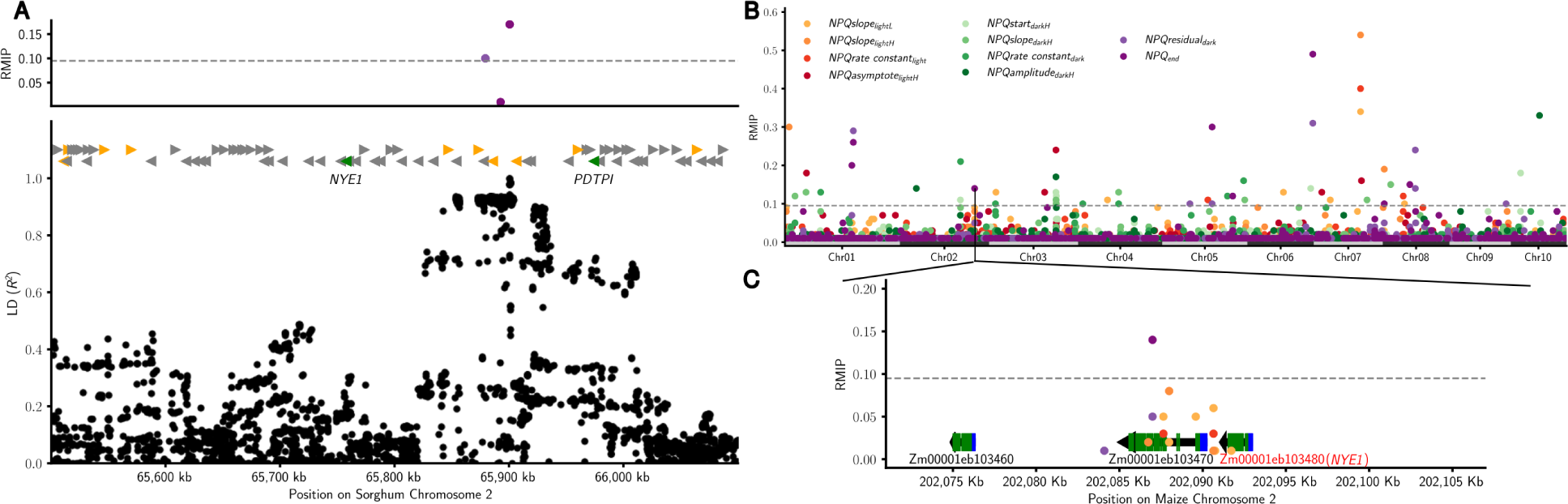
*NON-YELLOWING 1* (*NYE1*) is associated with variation in NPQ kinetics in sorghum and maize. **A)** Trait-associated SNPs, positions of candidate genes, and lineage disequilibrium for the LD defined mapping interval associated with hit 3 in Figure 2A and Supplementary Table S1: 65,500,078 to 66,099,985 bp on chromosome 2. This region contains two SNPs that pass the RMIP threshold Chr02:65,879,368 (*NPQresidual_dark_*, RMIP = 0.1) and Chr02:65,900,630 (*NPQ_end_*, RMIP = 0.17). The top panel shows trait-associated SNPs within this interval (colors correspond to traits given in the legend of panel B). Triangles indicate the position of annotated sorghum genes within the interval. Gray triangles indicate genes provisionally ruled out as candidates based on annotated function and/or lack of expression in sorghum leaf tissue. Yellow triangles indicate genes that could not be ruled out but lack clear links to chloroplasts, or photo-synthesis. Green triangles indicate the position of high-priority candidate genes based on functional annotation. Black dots indicate the LD of SNPs within this mapping interval relative to Chr02:65,879,368. **B)** RMIP results for a reanalysis of published trait data from 746 maize genotypes from the Wisconsin Diversity panel grown in Lincoln, Nebraska, in the summer of 2020 (Sahay *et al*. 2023) using new higher density genetic markers from resequencing (Grzybowski *et al*. 2023). **C)** Zoomed in view of the results from panel B for the region from 202,081 to 202,096 kb on maize chromosome 2. Colors of RMIP results correspond to the NPQ trait labels in panel B. Zm00001eb103480, the gene labeled in red is one of the two maize co-orthologs of *NYE1*, one of the green colored triangles genes in panel A. The gene adjacent to *NYE1*, Zm00001eb103470, is orthologous to a sorghum gene considered a low-priority candidate as it was expressed below 10 TPM in sorghum leaf tissue (Supplemental Data Set S4). Data used to produce this figure is given in Supplemental Data Sets S5A.

We reanalyzed previously published maize NPQ kinetics data (Sahay *et al*. 2023) using a recently published high-density marker set (Grzybowski *et al*. 2023) to prioritize candidate genes within the large LD interval present around the Chr02:65,879,368 and Chr02:65,900,630 markers in sorghum. This analysis identified 113 marker-trait associations with RMIP *≥* 0.1 (Figure 3B). One of these signals came from a genetic marker on maize chromosome 2 at position 202,086,988, 4.1 kb downstream of Zm00001eb103480. Zm00001eb103480 is one of two maize co-orthologs of Sobic.002G274800, one of the high-priority candidate genes above. Zm00001eb103480 and Sobic.002G274800 are both orthologs of *NYE1* (At4g22920), which encodes a chloroplast-localized protein involved in chlorophyll catabolism in *A. thaliana* (Park *et al*. 2007). The 6-kb interval from 202,086 to 202,092 kb on maize chromosome 2 included a cluster of additional marker-trait associations for three different methods of quantifying the rate of NPQ induction under high light: *NPQslope_lightH_*(aggregate RMIP=0.11, three markers), *NPQrate constant_light_*(aggregate RMIP=0.06, two markers, and *NPQslope_lightL_*(aggregate RMIP = 0.20, six markers) (Figure 3C). These three metrics estimate the rate of NPQ induction from hyperbolic, exponential, and linear functions, respectively. We also identified a statistically significant peak corresponding to the location of Zm00001eb103480 in a conventional mixed-linear-model-based GWAS for *NPQslope_lightL_* (Supplemental Figure S6).

Sorghum genotypes differing in their allele for the GWAS hit associated with *NYE1* also differed significantly in their values for (*NPQslope_lightH_*) and (*NPQ_end_*) (Figures 4A,B), consistent with the cluster of GWAS hits for these two traits associated with the maize ortholog of this gene (Figure 3C). Sorghum homozygous for the T allele exhibited significantly slower induction of NPQ (*NPQslope_lightH_*), lower steady state of NPQ induction (*NPQasymptote_lightH_*), lower maximum NPQ (*NPQ_max_*), and a smaller relaxation of NPQ after ten minutes in the dark (*NPQstart_darkH_*, *NPQamplitude_darkH_*, and *NPQ_end_*) than sorghum homozygous for the A allele (Figures 4A,B). Similarly, maize homozygous for TC allele had significantly lower induction of NPQ under high light conditions (*NPQslope_lightH_*) than did maize homozygous for the T-allele and smaller relaxation after 10 minutes in the dark (*NPQ_end_*) (Figures 4C,D). Two (*A. thaliana*) mutant lines carrying T-DNA insertions in *NYE1* (Figure 4E) exhibited largely consistent phenotypes with each other. Both lines reached lower maximum NPQ values under high light (*NPQasymptote_lightH_*, *NPQ_max_*, and *NPQstart_darkH_*) and relaxed more slowly in the dark than wild type (*NPQslope_darkH_*and *NPQamplitude_darkH_* (one of two lines)) (*P ≤* 0.04 for all listed comparisons) (Figure 4F,G,H). Unlike the differences observed between natural variants in sorghum and maize, no statistically significant differences in *NPQ_end_* were observed between *nye1* mutants and wildtype.

**Figure 4.**
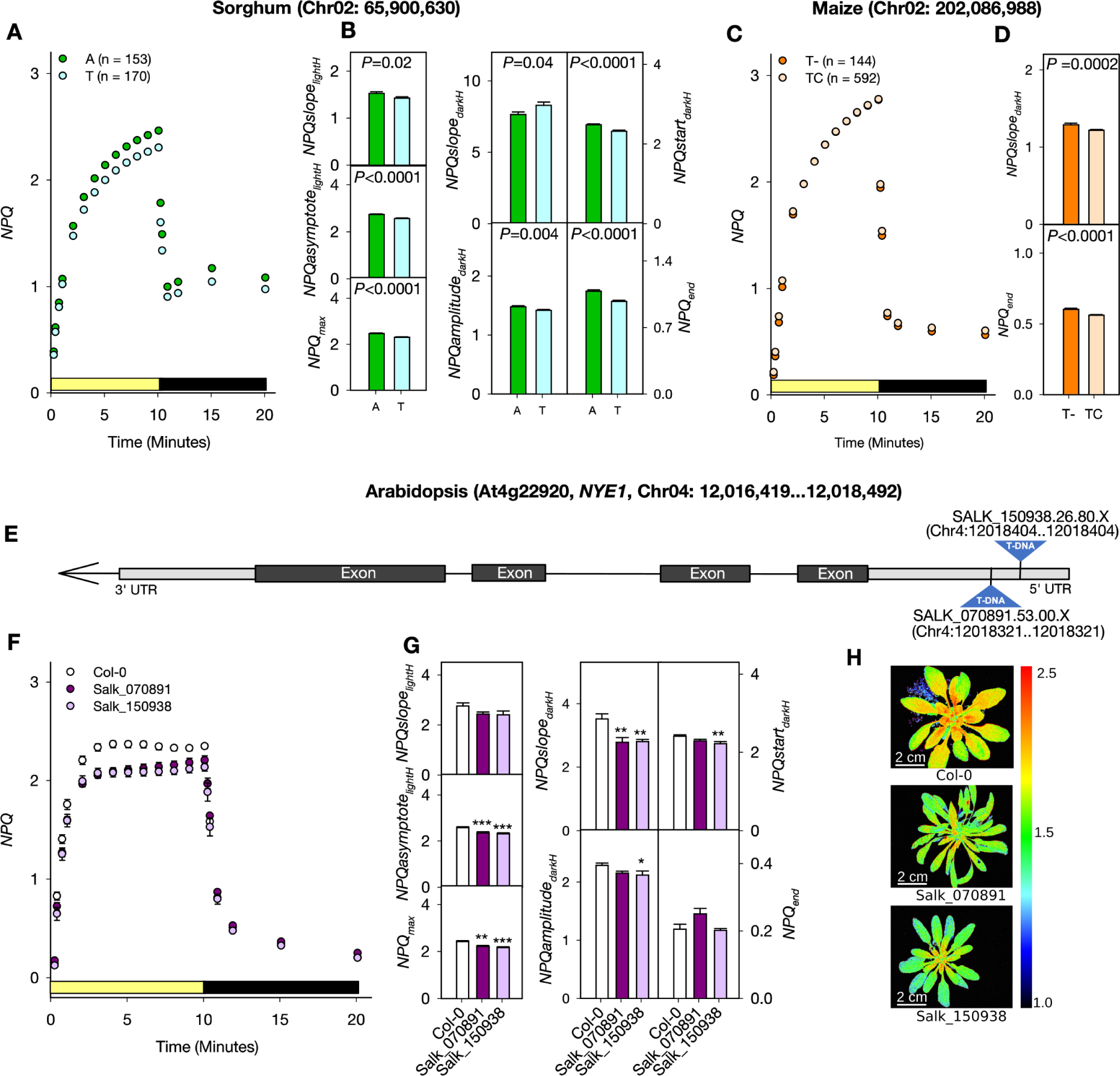
Nonphotochemical quenching (NPQ) phenotypes of *NON-YELLOWING 1* (*NYE1*) in sorghum, maize, and arabidopsis. **A)** Average pattern of NPQ induction and relaxation observed during a 10-minutes exposure to high-light condition (yellow bar) and relaxation observed during a subsequent 10-minutes dark treatment (black bar) for sorghum genotypes homozygous either the A or T allele at marker Chr02:65,900,630. Circles indicate median values, and standard error is indicated by vertical lines too small to be seen behind circles. **B)** Differences in a set of NPQ kinetic traits between sorghum homozygous for different alleles at position Chr02:65,900,630. Height of bars indicate mean values, error bars indicate standard error (n = 153 sorghum varieties for A allele and n = 170 sorghum varieties for T allele). **C-D)** Average pattern of NPQ kinetics and its traits as described in panel A, for maize homozygous for either the T-or TC allele of a single base InDeL located at position Chr02:202,086,988. Circles in panel C indicate median values, and the standard error is indicated by vertical lines too small to be seen behind circles. Bars in panel D indicate mean values and error bars standard error (n = 144 maize varieties for T-allele and n = 592 maize varieties for TC allele). In panel B and D, the *P*-values are the result of a test for significant differences between alleles (*P ≤* 0.05, two-tailed t-test). **E)** Structure of the *NYE1* (At4g22920) locus in (*A. thaliana*) and positions of the two T-DNA insertions used in this study. Dark gray boxes indicate protein-coding exons. Light gray boxes indicate untranslated regions. The arrow indicates the direction of transcription of *NYE1*. **F)** Pattern of NPQ induction and relaxation kinetics observed for wildtype *A. thaliana* (Col-0) and two *nye1* insertion lines (Salk_070891 and Salk_150938). **G)** Differences in a set of NPQ kinetic traits between Col-0 and *nye1* insertion lines. Bars indicate mean values and error bars standard error (n = 9 biological replicates for Col-0, n = 7 for Salk_070891, n = 6 for Salk_150938). Asterisks indicate significant differences between wildtype and mutants based on Dunnett’s two-way test: \**P ≤* 0.05; \*\**P ≤* 0.01; \*\*\**P ≤* 0.001. **H)** Chlorophyll fluorescence image of NPQ (Scale bar = 2cm) in Col-0, Salk_070891 and Salk_150938 after 10 minutes of high light illumination. Data used to produce this figure is given in Supplemental Data Sets S1, S6, S7 and S8.

Subsequent to successful mutant validation of the impact of *NYE1* on NPQ kinetics, we conducted a genome-wide search for maize and sorghum gene pairs adjacent to GWAS hits for the same NPQ kinetic phenotypes. We identified 10 maize-sorghum syntenic gene pairs in four genomic intervals that were adjacent to GWAS hits for traits measured as part of this study in 2020 (Figure 5). Six of these gene pairs corresponded to the *NYE1* locus itself. Two gene pairs on sorghum chromosome 9 and maize chromosome 6 were located in a window associated with *NPQstart_darkH_*in both maize and sorghum. These gene pairs are Sobic.009G204400–Zm00001eb293550 and Sobic.009G204601–Zm00001eb293530. A single gene pair on sorghum chromosome 2 (Sobic.002G125800) and maize chromosome 7 (Zm00001eb306410) was associated with the speed of NPQ induction (*NPQslope_lightL_*) in both species, with the two genes encoding a phosphoglucomutase. A single gene pair, Sobic.003G082800– Zm00001eb125070, on sorghum chromosome 3 and maize chromosome 3 was associated with *ϕPSIIamplitude_darkH_* and *ϕPSII_end_* in both species (the protocol employed in both this study and the previous maize study also enabled the quantification of the quantum yield of photosystem II (*ϕ*PSII) (Sahay *et al*. 2023)). The single most consistently identified signal for *ϕPSIIamplitude_darkH_* (RMIP=0.44) in sorghum was located near Sobic.005G075900, the ortholog of a largely uncharacterized *A. thaliana* gene that is primarily expressed in leaves and encodes a cysteine/histidine-rich C1 domain family protein (At2g16050). The maize ortholog of this sorghum gene, Zm00001eb168930, was also associated with GWAS hits for multiple *ϕ*PSII kinetic traits and is primarily expressed in mature leaves and roots (Stelpflug *et al*. 2016).

**Figure 5.**
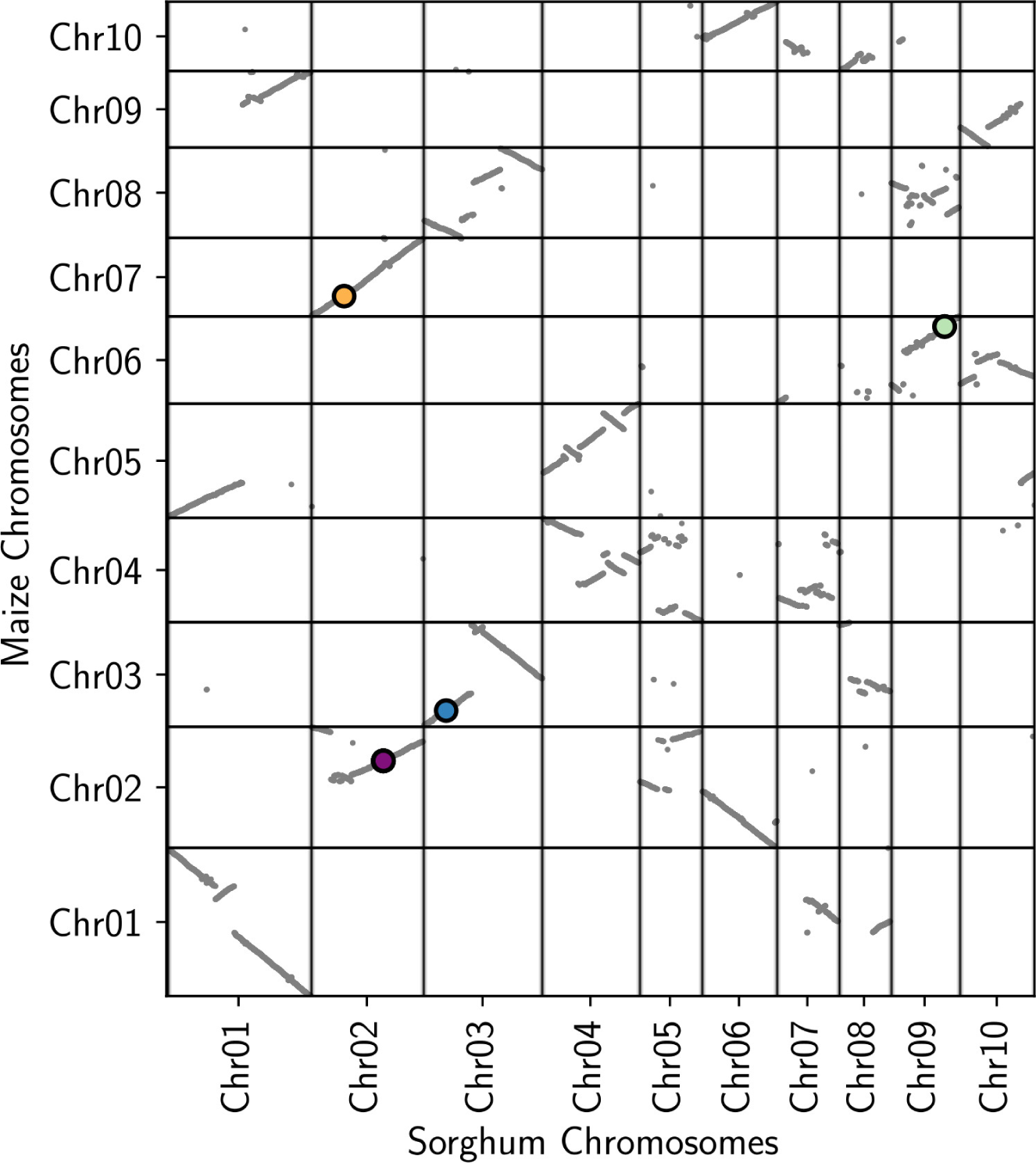
Syntenic GWAS hits for photosynthetic phenotypes identified across maize (2020) and sorghum (LN2020). The x- and y-axes indicate annotated gene order along the genomes of sorghum (BTx623 v3) and maize (B73 RefGen v5), respectively. Gray dots mark the locations of maize-sorghum syntenic orthologous gene pairs. Large colored circles with black outlines indicate the positions of syntenic orthologous gene pairs in which the maize gene copy is no more than 100 kb from a GWAS hit for one or more traits identified with an RMIP *≥* 0.05 ("suggestive") and the sorghum gene copy is no more than 300 kb from a GWAS hit for the same trait or traits identified at the same RMIP *≥* 0.05 ("suggestive") cutoff. The filled colors of each circle (orange: *NPQslope_lightL_*, purple: *NPQ_end_*, blue: *ϕPSIIamplitude_darkH_*, and green: *NPQstart_darkH_*) indicate the identity of the shared trait between maize and sorghum orthologs. When multiple shared traits were identified, the trait with the greatest average RMIP across the two species was used to select the color shown.

## Discussion

In this study, we quantified various parameters related to NPQ kinetics for 339 sorghum genotypes grown under both low-nitrogen and high-nitrogen conditions in 2020 and 2021 and performed GWAS using high-density genetic markers from resequencing data (Boatwright *et al*. 2022). We detected substantial heritable variation in NPQ kinetics phenotypes among sorghum genotypes, although the proportion of variance attributable to genetics was modestly lower than that observed in a maize association panel for the same traits (Table 2). The GWAS conducted in the environment with the most heritable measurements identified 20 unique marker-trait associations for five different NPQ kinetics traits across the sorghum genome. The most consistently identified signal was associated with the sole sorghum ortholog of an (*A. thaliana*) gene implicated in plastid–nucleus signaling and tolerance to stresses likely to produce ROS (Figure 2). However, in many cases, the mapping windows for GWAS hits in sorghum were too broad to identify a single high-confidence candidate gene (Figure 3A). Reanalyzing published NPQ kinetics data from maize using a new and higher density marker set (Grzybowski *et al*. 2023; Sahay *et al*. 2023) led to the identification of Mendel’s green pea gene *NON-YELLOWING 1* (*NYE1*) as a gene associated with variation in the same NPQ kinetics phenotype in both maize and sorghum populations (Figure 3B,C). We validated this association via knockout phenotypes in (*A. thaliana*) (Figure 4E-H). Given the success of this multi-species GWAS approach, we conducted a systematic analysis of all GWAS hits identified in maize and sorghum and identified three additional syntenic GWAS hits for future characterization (Figure 5). These gene pairs with similar positions and associations with the same trait across both species provide evidence of a conserved functional role of proteins in controlling NPQ.

The most consistently identified signal in our sorghum NPQ kinetics GWAS flagged the sorghum ortholog of *SUPPRESSOR OF VARIRGATION3* (*SVR3*) which encodes a putative Type A translation elongation factor localized to the chloroplast (Liu *et al*. 2010). *SVR3* was initially identified as a suppressor of the variegation2 mutant phenotype and caused bleaching in the light at low temperature (Liu *et al*. 2010). Another allele of the same gene was isolated based on resistance to norflurazon and accumulated more carotenoids than wildtype (Saini *et al*. 2011). Two loss-of-function alleles of *SVR3* accumulated less photosystem II reaction center protein D1 than wildtype plants under control conditions (Liu *et al*. 2010). Mutations of *SVR3* fail to produce functional chloroplasts at low temperatures (Liu *et al*. 2010; Saini *et al*. 2011). Chilling stress slows the enzymes of the Calvin-Benson-Bassham cycle more than the enzymes of the light-dependent reactions of photosynthesis, resulting in the harvesting of excess energy that cannot be consumed by the Calvin-Benson-Bassham cycle and produces ROS (Wise 1995), creating some of the same energy balance challenges stemming from rapid increases in light intensity. This pattern of change in NPQ kinetics observed in sorghum – more rapid induction and higher maximum values of NPQ under high-light treatment, but slower and less complete relaxation of NPQ in the dark (Figure 2C) – is consistent with a larger pool of NPQ-related carotenoids increasing the magnitude of qZ.

A combined GWAS across maize and sorghum identified orthologs in each species of *NYE1* the gene responsible for the segregating cotyledon color phenotype used by Gregor Mendel in determining the laws of genetics (Armstead *et al*. 2007; Sato *et al*. 2007). Orthologs of this gene have been cloned in many species, as their loss-of-function mutants frequently exhibit delayed or lack of leaf yellowing during senescence (Armstead *et al*. 2007; Jiang *et al*. 2007; Park *et al*. 2007; Ren *et al*. 2007). The *NON-YELLOWING1* genes encode magnesium-dechelatase, responsible for catalyzing the first committed step of chlorophyll breakdown, the removal of the Mg^+2^ from Chlorophyll *a* (Shimoda *et al*. 2016). In *A. thaliana*, *NYE1* expression is upregulated in NPQ-deficient mutants under high-light conditions (Alboresi *et al*. 2011); in rice, natural variation in the expression of the *NYE1* ortholog *STAY-GREEN* (*SGR*) is associated with differences in photosynthetic productivity and grain yield (Shin *et al*. 2020). The alleles of the *NYE1* GWAS hits in maize and sorghum that were associated with faster induction of NPQ under high light conditions were also associated with less complete relaxation of NPQ in the dark subsequent to high light treatment (Figure 4).

At a population level, the initial speed of NPQ induction (*NPQslope_lightH_*) was only modestly correlated with either the maximum level of NPQ under high light treatment (*NPQasymptote_lightH_*, R^2^ = 0.01), the speed of relaxation in the dark (*NPQslope_darkH_*, R^2^ = 0.00), or maximum degree of relaxation (*NPQ_end_*, R^2^ = 0.04) (Supplemental Figure S7). However, the maximum level of NPQ under high-light treatment was substantially correlated with the final degree of NPQ relaxation in the dark (R^2^ = 0.64), suggesting a potential trade-off between the capacity to adapt to high-light conditions and the ability to optimize photosynthetic productivity under lower light conditions rapidly (Supplemental Figure S7). Here, we identified individual genetic variants via GWAS for individual traits describing different features of the kinetics of NPQ induction and relaxation. However, in some cases, GWAS hits for multiple NPQ kinetics phenotypes clustered on the sorghum and maize genomes (Figures 2A, 3B). In addition, analysis of the overall pattern of NPQ kinetics in genotypes carrying different alleles of trait-associated markers identified by GWAS (Figure 2C, 2D, 4A-D) and mutant alleles (Figure 4F 4G) suggest that functional variation in a single locus can alter multiple aspects of NPQ kinetics. As a result, it may be necessary to characterize larger numbers of genes to identify a specific genetic intervention or intervention that can alter the kinetics of NPQ in a desired fashion. Fortunately, the consequences of functional variation in orthologous genes appear to be conserved across even distantly related species, as reported previously for five cases in maize and *A. thaliana* (Sahay *et al*. 2023) and described here for *NYE1* in maize, sorghum, and *A. thaliana*, allowing the results of gene characterization to be shared across different crops and genetic models.

Conservation of phenotypes across orthologous genes has been widely reported among the grasses, including the domestication gene *Shattering1* (Lin *et al*. 2012) and the flowering time regulator *Heading date1* (Liu *et al*. 2015). Conservation of function has also been reported in the cloned genes via classical forward genetics as in the case of sorghum *Dwarf 3* (*dw3*) and maize *Brachytic2* (*br2*), which encode orthologous modulators of polar auxin transport involved in stem elongation (Multani *et al*. 2003). Maize has served as a genetic model for more than a century, with hundreds of genes subjected to individual investigation and characterization (Schnable and Freeling 2011). However, forward genetic characterization in sorghum has been more limited (Boyles *et al*. 2019). GWAS provides an alternative approach to link genes to functions, but the slower decay of LD in the sorghum genome frequently results in mapping intervals that are too large to identify a single candidate gene confidently. This was the case with the GWAS hit identified at Chr02:65,900,630 in this study. The approach we took to identify *NYE1* integrated GWAS results for the same phenotype across maize and sorghum. This approach builds upon prior findings that demonstrated the conservation of a number of flowering time QTLs across maize, sorghum, and foxtail millet (*Setaria italica*) (Mauro-Herrera *et al*. 2013) and root phenotype associated GWAS hits across maize and sorghum (Zheng *et al*. 2020). We suspect the approach employed here may also be applicable to other traits and pairs of related species.

## Materials and methods

### Composition and Design of Sorghum Field Experiment

The field data presented here were collected from two experiments conducted in the summers of 2020 and 2021 at the University of Nebraska-Lincoln’s Havelock Farm. In both years plots consisted of one 2.3-m row of plants from a single genotype planted with a spacing of 0.76 m between parallel and sequential rows and approximately 3.8 cm in spacing between sequential plants in the same row. In both years, no supplemental nitrogen was applied to low-nitrogen (LN) treatment blocks and a target of 80 pounds of N per acre (89.7 kilograms/hectare) was applied to high-nitrogen (HN) blocks. The 2020 experiment was conducted in a field located at N*^◦^* 40.861, W*^◦^* 96.598, and the 2021 experiment was conducted in a field located at N*^◦^* 40.858, W*^◦^* 96.596. The 2020 experiment consisted of 347 genotypes drawn from the sorghum association panel plus Tx430 (Casa *et al*. 2008). A total of four blocks were planted, each consisting of 390 plots with one entry per genotype, plus BTx623 as a repeated check. Two blocks were grown under HN treatment and two under LN treatment. The 2021 experiment consisted of 347 genotypes replicated in two blocks, each consisting of 390 plots including the replicated check, under LN conditions as in 2020 and 911 sorghum genotypes in two blocks, each consisting of 966 plots including the replicated check under HN. The larger population in HN treatment resulted from the inclusion of sorghum genotypes from the sorghum diversity panel (Griebel *et al*. 2021) in addition to the previously characterized sorghum association panel lines. The 2020 field experiment was planted on June 8^th^, and the 2021 field experiment was planted on May 25^th^. The 2020 field experiment has also been previously described (Grzybowski *et al*. 2022).

### Quantifying NPQ and Deriving NPQ Kinetics Traits

The leaf disks employed for NPQ measurements were collected from the fields between July 29^th^ and August 5^th^ in 2020 and between July 21^st^ and July 30^th^ in 2021, with collection occurring between 16:00 and 18:30 hours. Leaf disks were collected and NPQ traits were profiled from leaf disks following the method described previously (Sahay *et al*. 2023). Briefly, for each plot, three plants were randomly selected for sampling, avoiding edge plants when possible. For each plant, one 0.32-cm^2^ leaf disk was collected from the youngest fully expanded leaf, avoiding the midrib. Leaf disks were placed with their adaxial surfaces facing down into 96-well plates, with wet sponges added onto their abaxial surfaces to prevent drying and secure disc position. Plates were inverted (i.e., adaxial face up) and incubated overnight in the dark prior to phenotyping. The following day, plate imaging was performed using a modulated chlorophyll fluorescence imager (FluorCam FC 800-C, Photon Systems Instruments, Drasov, Czech Republic). Leaf disks were subjected to an illumination treatment of 10 minutes at 2,000 *µ*mol m^-2^s^-1^ light (1,000 *µ*mol m^-2^s^-1^ of red-orange light with *λ*_max_ = 617 nm and 1,000 *µ*mol m^-2^s^-1^ of a 6,500K light source) followed by 10 minutes of dark treatment with saturating flashes of 3,200 *µ*mol m^-2^s^-1^ (provided by a cool white 6,500K light) for a duration of 800 ms. The minimum (*F_o_*) and maximal (*F_m_*) chlorophyll fluorescence values were measured from the dark-adapted leaves, followed by steady-state fluorescence (*F_s_*) and maximum fluorescence measured under illuminated conditions (*F_m_^’^*) at the following intervals (in seconds): 15, 30, 30, 60, 60, 60, 60, 60, 60, 60, 60, 60, 60, 9, 15, 30, 60, 180, and 300. At each time point, NPQ was estimated via the Stern-Volmer quenching model (Bilger and Björkman 1994).

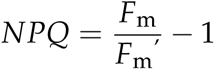

The measured NPQ represents the two fastest components: energy-dependent quenching (qE) and modestly slower zeaxanthin-dependent quenching (qZ). NPQ values calculated at each time point were fitted to hyperbola and exponential equations in MATLAB (Matlab R2019b; MathWorks, Natick, MA, USA) to extract parameters reflecting the rate, amplitude, and steady state of the NPQ kinetics curve (Table 1) using the equations described in (Sahay *et al*. 2023). In both years, plots with no healthy plants present were excluded from sampling and analysis. In 2020, 1,550 of 1,560 plots were sampled. In 2021 2,667 of 2,712 plots were sampled.

### Data Integrity and Quality Control Procedure

Individual leaf disks were subjected to a two-stage quality control and outlier removal process with the first stage based on maximum efficiency of PSII (*F_v_/F_m_*) and goodness of fit for hyperbolic equations fit to the measured NPQ and the second stage based on visual examination of the distribution of each NPQ kinetic trait to identify extreme values. After quality control and outlier removal, data was present for 348 sorghum genotypes in both treatments across both years (Supplemental Data Set S1). Within-year/within-treatment Best Linear Unbiased Estimators (BLUEs) were calculated by fitting a mixed linear model using the *lme4* package (Bates *et al*. 2015) in R v.4.1.0 (R Core Team 2020) with the equation *y_i_* = *µ* + *Genotype_i_* + *Row_j_* + *Block_k_* + *Plate_l_* + *error_ijkl_* where, *y_i_* is the mean trait of interest in the *i^th^* genotype planted in the *j^th^* row and *k^th^* block, and placed in *l^th^* 96-welll plate during NPQ quantification, *µ* is the overall mean of the population, *Genotype_i_* is fixed effect of genotype *i*, *Row_j_* is random effect of row *j*, *Block_k_* is random effect of block *j*, *Plate_l_* is random effect of 96-well plate *l*, and *error_ijkl_* is the residual. Visual examination of BLUE distributions identified nine sorghum genotypes (*∼*2.6% of the total population) with extreme values for one or more NPQ traits. BLUEs for 339 sorghum genotypes (Supplemental Data Set S2) were employed for genome-wide association studies.

The variance explained by individual factors within the model was used to estimate within-year/within-treatment heritability using the following equation:

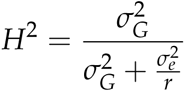

Where 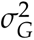 was the total variance explained by the genotype factor in each mixed linear model and 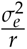 is the total residual variance in each mixed linear model divided by the number of replicates of each genotype within the same treatment and year.

### Quantitative Genetic Analyses

A published set of 43 million segregating single nucleotide polymorphisms (SNPs), structural variants, and insertion/deletions (InDels) derived from whole–genome resequencing of the sorghum association panel (Boatwright *et al*. 2022) was filtered using VCFtools v.0.1 (Danecek *et al*. 2011) and bcftools v.1.17 (Danecek *et al*. 2021) retaining only biallelic SNP markers with a minor allele frequency (MAF) >0.05 and a frequency of heterozygous genotype calls <0.05 among the final set of 339 sorghum genotypes. These filtering criteria produced a final set of 4,457,229 SNP markers.

Marker-trait associations in sorghum were considered significant when identified in at least 10 of 100 resampling analyses conducted with FarmCPU as implemented in rMVP (Liu *et al*. 2016; Yin *et al*. 2021). In each iteration, a random 10% of sorghum genotypes were masked, and a separate FarmCPU analysis was conducted, keeping the set of markers significantly above a threshold of 5.107 *×* 10^-8^. The significance threshold employed corresponds to 0.05 divided by the approximately 978,957 effective/independent number of markers in the filtered biallelic SNP dataset as estimated by GEC v.0.2 (Li *et al*. 2012).

Marker-trait association in maize was conducted using a published set of maize NPQ kinetics phenotypes (Sahay *et al*. 2023). FarmCPU resampling was conducted as described above, with a set of 16,634,049 markers representing the subset of the 46 million high-confidence markers identified previously (Grzybowski *et al*. 2023) with a minor allele frequency >5% and significant threshold 3*×*10*^−^*^9^.

Linkage disequilibrium (LD) calculations were performed using plink v1.90; (Purcell 2012). Windows of interest were defined by first calculating LD between the marker identified by FarmCPU resampling and all SNPs within an interval of approximately 10 Mb and then identifying the first and last SNPs within the interval that exhibited linkage disequilibrium with the GWAS hit above the cutoff used to define the window.

### Quantification of NPQ in *nye1* Mutants

The seeds of *A. thaliana* T-DNA insertion *nye1* mutants (Salk_070891 and Salk_150938) and the corresponding Columbia (Col-0) wildtype control (CS60000) were obtained from the Arabidopsis Biological Resource Center (ABRC) at Ohio State University (Alonso *et al*. 2003). Lines were selfed and homozygous progeny was obtained. Homozygosity for *nye1* mutant plants was confirmed by PCR using the T-DNA-specific primer LBb1.3 and gene-specific left primer and right primer for Salk_070891 and Salk_150938 lines (Supplemental Table S2) which were designed using the Salk T-DNA primer design tool (http://signal.Salk.edu/tdnaprimers.2.html). The seeds of homozygous T-DNA mutant lines and their corresponding wildtype were stratified at 4*^◦^*C for four days in the dark, sown in 3.5 inch *×* 3.5 inch (8.89 *×* 8.89 centimeter) pots (SQN03500B66, Hummert International, Earth City, MO, USA) filled with soil-less potting mix (1220338; BM2 Germination and Propagation Mix; Berger, SaintModeste, Canada) at 21*^◦^*C day/18*^◦^*C night with a 10-h-light/14-h-dark photoperiod (200 *µ*mol m^-2^ s^-1^) and 60% relative humidity in a reach-in growth chamber (AR-66L2; Percival, Perry, IA, USA). One week after germination, seedlings were thinned to one per pot; plants were watered three times a week with 150 ppm liquid fertilizer (Peter’s 201020 general purpose fertilizer, 25#, Peters Inc., Allentown, PA, USA). Plants were repositioned three times a week at random locations in the chamber.

NPQ kinetics were measured and parameterized as above, except with a few modifications described below. Four-week-old whole plants were imaged after 20 minutes of dark adaptation, with saturating pulses of 2,400 *µ*mol m^-2^ s^-1^ and actinic light of 1,000 *µ*mol m^-2^ s^-1^ (a combination of 500 *µ*mol m^-2^ s^-1^ of a red-orange light with *λ*_max_=617nm and 500 *µ*mol m^-2^ s^-1^ of a cool white 6500K light) were used. An area of 270–280 pixels was chosen manually from a portion of three of the youngest, fully expanded leaves facing the uniform illumination and used for NPQ quantification and analysis.

### Comparative GWAS Analysis

The coding sequences of annotated maize genes from B73 RefGen_V5 (Hufford *et al*. 2021) and annotated sorghum genes from the BTx623 V3 (McCormick *et al*. 2018) reference genomes were downloaded from Phytozome, selecting the "primaryTranscriptOnly" files for each species (Goodstein *et al*. 2012). Syntenic orthologs were identified by 1) identifying genes with similar coding sequences between the two species using LASTZ with a seed of match12, a minimum identity of 70% and minimum coverage of 50% (Harris 2007), 2) identifying syntenic blocks of homologous genes using QuotaAlign with a quota of 1:2 for sorghum:maize (Tang *et al*. 2011), reflecting the whole-genome duplication that occurred in maize after the split of the maize and sorghum lineages; and 3) polishing the raw QuotaAlign gene pairs via the Zhang et al. method (Zhang *et al*. 2017).

### Statistical Data Analysis of Insertion Mutant Phenotypes

Statistical analysis of NPQ kinetics measurements in mutants was performed using SAS (version 9.4, SAS Institute Inc., Cary, NC, USA). Normality of distributions was evaluated using the Shapiro–Wilk test and the homogeneity of variance in data was evaluated using the Brown–Forsythe method. When the assumption of either normality or homogeneity of variance were violated as determined by the above methods data were log transformed prior to testing for significant differences between genotypes. To evaluate potential differences between mutant and wild type plants, one-way ANOVAs (*α* = 0.05) were performed followed by Dunnett’s test to address the multiple comparison problem of multiple insertion lines being compared to a common wildtype control.

## Supporting information

Archive of supplementary data files

## Data Availability

NPQ data collected from sorghum are provided as Supplementary Data Sets associated with this paper. Raw measurements of NPQ parameters for each leaf disk at each time point are provided for individual leaf disks (Supplemental Data Set S1). Within year within treatment BLUEs for NPQ kinetic phenotypes are provided as Supplementary Data Set S2. The genetic marker data employed in this study has been previously published (Boatwright *et al*. 2022; Grzybowski *et al*. 2023). The sorghum genetic marker data is available from the European Variant Archive (PRJEB51985) and the maize genetic marker data is available from Dryad: https://doi.org/10.5061/dryad.bnzs7h4f1. The set of FarmCPU GWAS hits identified for all traits described in this study for sorghum are provided as Supplemental Data Set S3. Annotated sorghum genes within the mapping interval for each GWAS hit presented in Supplemental Table S1 are provided in Supplemental Data Set S4. The set of FarmCPU GWAS hits identified in maize using resequenced marker data are provided as Supplemental Data set S5. Data of genotype calls across the sorghum association panel for representative GWAS hits (Supplemental Data Set S6) and across the Wisconsin Diversity maize panel for GWAS hit corresponding to *NYE1* (Supplemental Data Set S7) are provided. Raw data of NPQ values for arabidopsis mutants are provided as Supplemental Data Set S8.

## Acknowledgments

The authors thank Mackenzie Zwiener, Brandi Sigmon, and Christine Smith for their leadership on field design and field experiment execution in 2020 and 2021. The authors thank Aldi Airori, Grace Carey, Sierra Conway, Elijah Frost, Remy Hirwa, Alliance Igiraneza, Luke Micek, Prince Ngiruwonsanga, Nathaniel Pester, Isabel Sigmon, Isaac Stevens, Lou Townsend, Daniella Norah Tumusiine, and Leighton Wheeler, for assistance with planting, field maintenance, and field data collection. The authors thank Annie Nelson for assistance in plate imaging and Eleanor Browne, Quinton Browne, Rachel Gerdes, Eleana Kang, Quinn Kimbell, Ryenne Leising, Bailey McClean, Pascaline Nyonshuti, and Hayley Ramirez for assistance in the collection of plant materials from the field. The authors thank Harshita Mangal for assistance in data analysis.

## Author contributions

JCS, MG, and KG conceived the project. SS collected the data. SS, NS, and MG analyzed the data. HMS and RVM conducted additional analyses. NS, JCS, and SS designed figures and drafted the manuscript. MG designed additional figures. All authors read, edited, and approved the final manuscript.

## Funding

This project was supported by the U.S. Department of Energy, Grants no. DE-SC0020355 and DE-SC0023138, the National Science Foundation under grant OIA-1826781, USDA-NIFA under the AI Institute: for Resilient Agriculture, Award No. 2021-67021-35329 and the Foundation for Food and Agriculture Research Award No. 602757 to JCS. This project was supported by the National Science Foundation under awards OIA-1557417 (CRRI and ESCoR FIRST) and IOS-2142993 (CAREER) and the University of Nebraska Lincoln under a Laymen award to KG. HMD supported by a FAPESP award, (support # 2022/16208-9).

## Conflicts of interest

James C. Schnable has equity interests in Data2Bio, LLC; Dryland Genetics LLC; and EnGeniousAg LLC. He is a member of the scientific advisory board of GeneSeek and currently serves as a guest editor for The Plant Cell. The authors declare no other competing interests.

## Supplemental data

The following materials are available in the online version of this article.

- **Supplementary Figure S1:** Diversity of NPQ induction and relaxation patterns observed in the sorghum association panel.
- **Supplementary Figure S2:** Distributions of NPQ trait across years and treatments.
- **Supplementary Figure S3:** Correlations of NPQ kinetics traits presented in Figure 1 across years and treatments.
- **Supplementary Figure S4:** Results of GWAS for NPQ kinetic traits measured for sorghum genotypes in two treatments over two years.
- **Supplementary Figure S5:** Zoomed in view of the Sobic.004G128900/*SVR3* candidate gene and LD window.
- **Supplementary Figure S6:** Conventional MLM-based GWAS identifies signals at the genomic interval corresponding to Zm00001eb10348 (*NYE1*).
- **Supplementary Figure S7:** Correlation among hyperbola derived NPQ trait in maize.
- **Supplemental Table S1:** Locations and mapping intervals for significant GWAS hits associated with NPQ relaxation in dark traits under low nitrogen treatment in 2020.
- **Supplemental Table S2:** Names and sequences of primers used for PCR to verify homozygosity of *A. thaliana* mutants in *NYE1* (At4g22920) gene.
- **Supplementary Data Set S1:** NPQ and *ϕ*PSII values at each time point during light and dark treatment for each sorghum leaf disk collected in 2020 and 2021.
- **Supplementary Data Set S2:** BLUE values for each sorghum genotype in 2020 and 2021 under HN and LN treatments. NA indicates values that were excluded based on analysis of trait distributions prior to GWAS.
- **Supplementary Data Set S3:** Results of resampling-based GWAS in sorghum. Includes analyses conducted in both HN and LN treatments in 2020 an 2021.
- **Supplementary Data Set S4:** Set of annotated sorghum genes present within the mapping intervals for each GWAS hit described in Supplemental Table S1.
- **Supplementary Data Set S5:** Results of resampling-based GWAS in maize conducted using new higher density marker data. Includes results of separate analyses conducted using published trait data from 2020 and 2021.
- **Supplementary Data Set S6:** Genotype calls across the sorghum association panel for trait-associated markers described in Supplemental Table S1. Heterozygous calls were converted to missing data.
- **Supplementary Data Set S7:** Genotype calls across the Wisconsin Diversity maize panel for Chr2:2,286,988 Allele information for GWAS hit corresponding to *NYE1* in maize. Empty cells were heterozygous alleles removed, as inbreds are not expected to have heterozygous alleles.
- **Supplementary Data Set S8:** NPQ values at each time point during light and dark treatment for each wild type and *nye1* plant for which data is reported in Figure 4.

---

**Figure S1.**
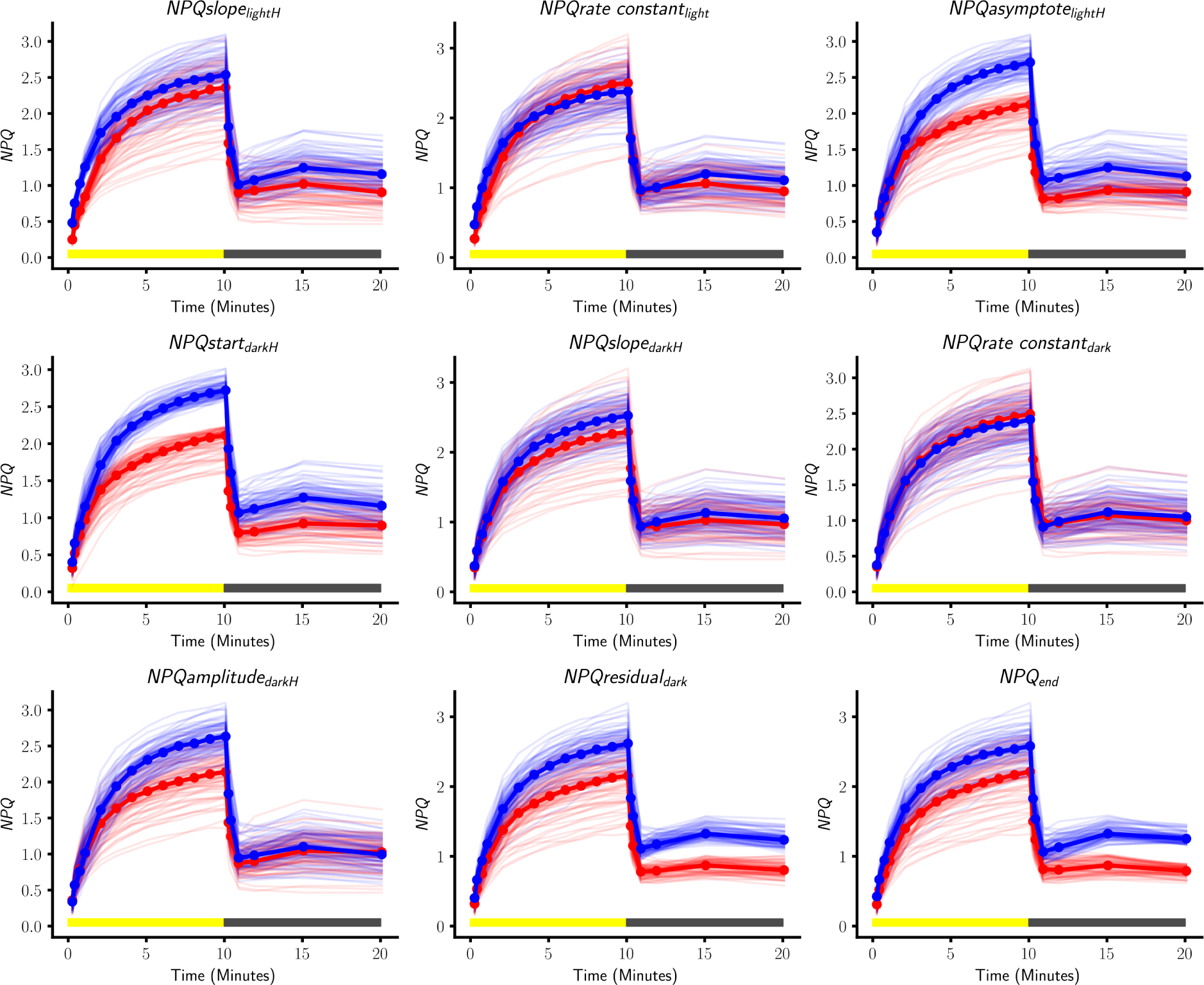
Diversity of NPQ induction and relaxation patterns observed in the sorghum association panel. Patterns of NPQ induction observed for individual genotypes in the sorghum association panel grown under low-nitrogen conditions in the summer of 2020. Each plot corresponds to one of nine traits calculated from the NPQ response curves. Solid red and blue lines indicate median NPQ responses among all genotypes ranking in the 2^nd^-25^th^ and 75^th^-98^th^ percentiles, respectively. Light red lines indicate NPQ response curves for sorghum genotypes between the 2^nd^ and 25^th^ percentile within the population for the given NPQ trait. Light blue lines indicate NPQ response curves for sorghum genotypes between the 75^th^ and 98^th^ percentile within the population for the given NPQ trait. Data used to produce this figure is given in Supplemental Data Set S1A.

**Figure S2.**
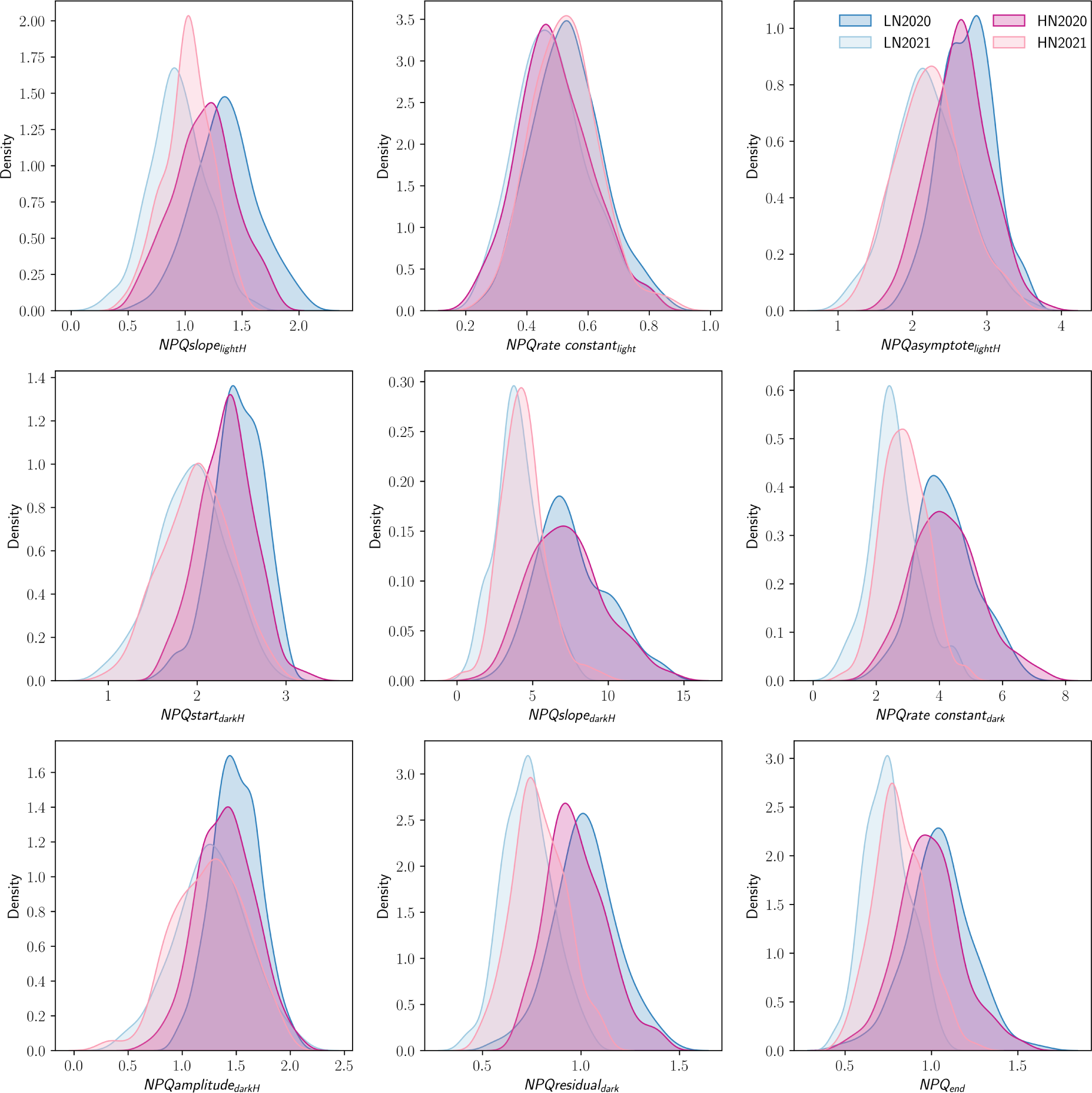
Distributions of NPQ trait across years and treatments. One-dimensional kernel density estimates of the distributions of best unbiased linear estimators + trait means calculated for sorghum genotypes under low-nitrogen conditions and nitrogen-sufficient conditions in 2020 and 2021. Within year paired t-test between treatments: *NPQslope_lightH_*; (*P* <0.0001 for both years, but with opposite effects of nitrogen across years), *NPQrate constant_light_*; (*P* <0.0001 for both years), *NPQasymptote_lightH_*; (*P* = 0.65 (2020), *P*<0.001 (2021)), *NPQstart_darkH_*; (*P* <0.001 (2020), *P* = 0.03 (2021), but with opposite effects of nitrogen across years), *NPQslope_darkH_*; (*P* = 0.04 (2020), *P* <0.001 (2021), but with opposite effects of nitrogen across years), *NPQrate constant_dark_*; (*P* = 0.96 (2020), *P* <0.0001 (2021)), *NPQamplitude_darkH_*; (*P* <0.0001 (2020), *P* = 0.44 (2021)), *NPQresidual_dark_*; (*P* <0.0001 for both years, but with opposite effects of nitrogen across years), *NPQ_end_*; (*P* <0.0001 for both years, but with opposite effects of nitrogen across years). Data used to produce this figure is given in Supplemental Data Set S2.

**Figure S3.**
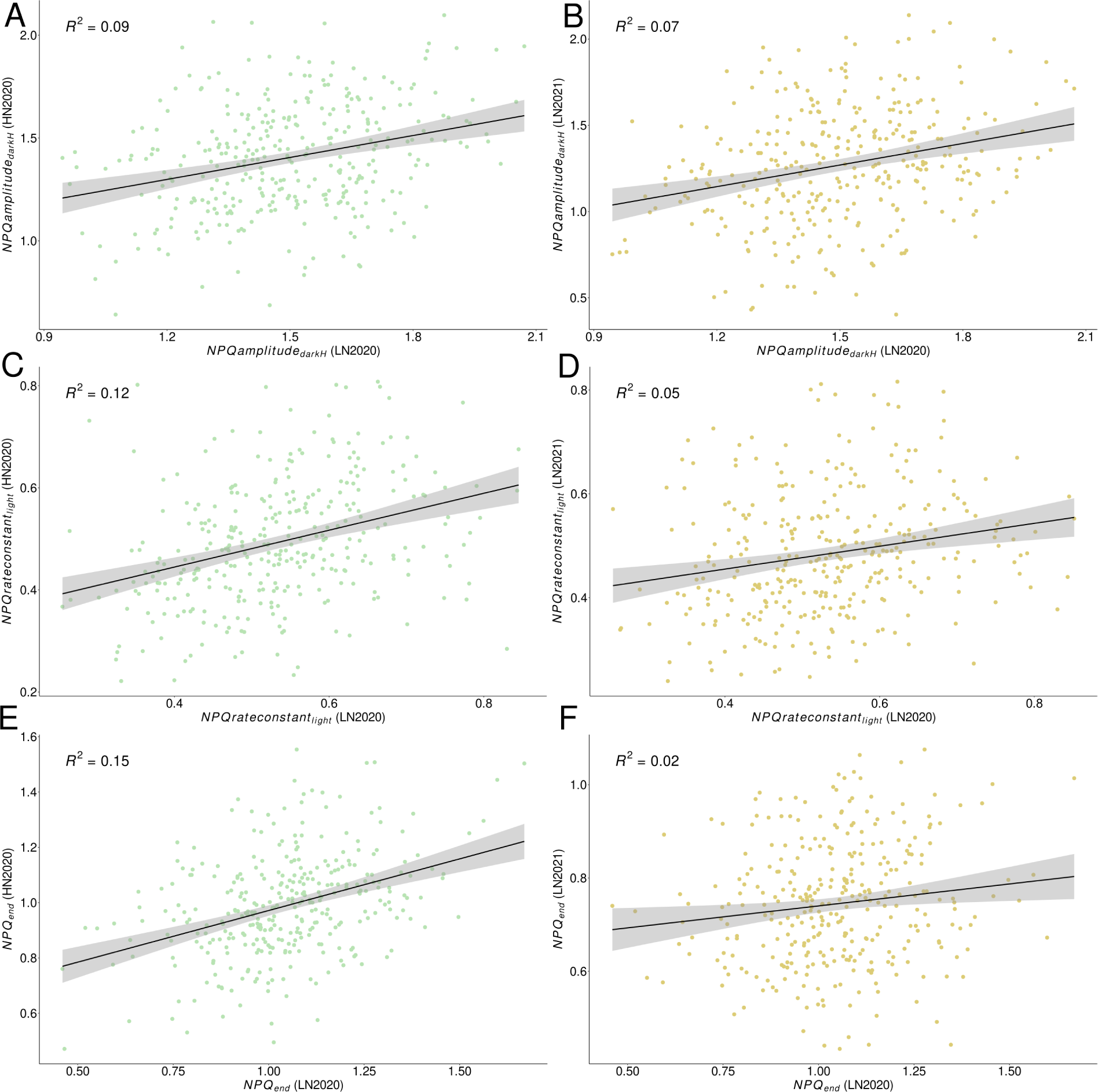
Correlations of NPQ kinetics traits presented in. **Figure 1 across years and treatments. A** Correlation between *NPQamplitude_darkH_* BLUEs for the same sorghum genotypes under HN and LN treatments in 2020. **B** Correlation between *NPQamplitude_darkH_* BLUEs for the same sorghum genotypes under LN treatment in 2020 and 2021. **C** Correlation between *NPQrate constant_dark_* BLUEs for the same sorghum genotypes under HN and LN treatments in 2020. **D** Correlation between *NPQrate constant_dark_* BLUEs for the same sorghum genotypes under LN treatment in 2020 and 2021. **E** Correlation between *NPQ_end_* BLUEs for the same sorghum genotypes under HN and LN treatments in 2020. **F** Correlation between *NPQ_end_* BLUEs for the same sorghum genotypes under LN treatment in 2020 and 2021. Data used to produce this figure is given in Supplemental Data Set S2.

**Figure S4.**
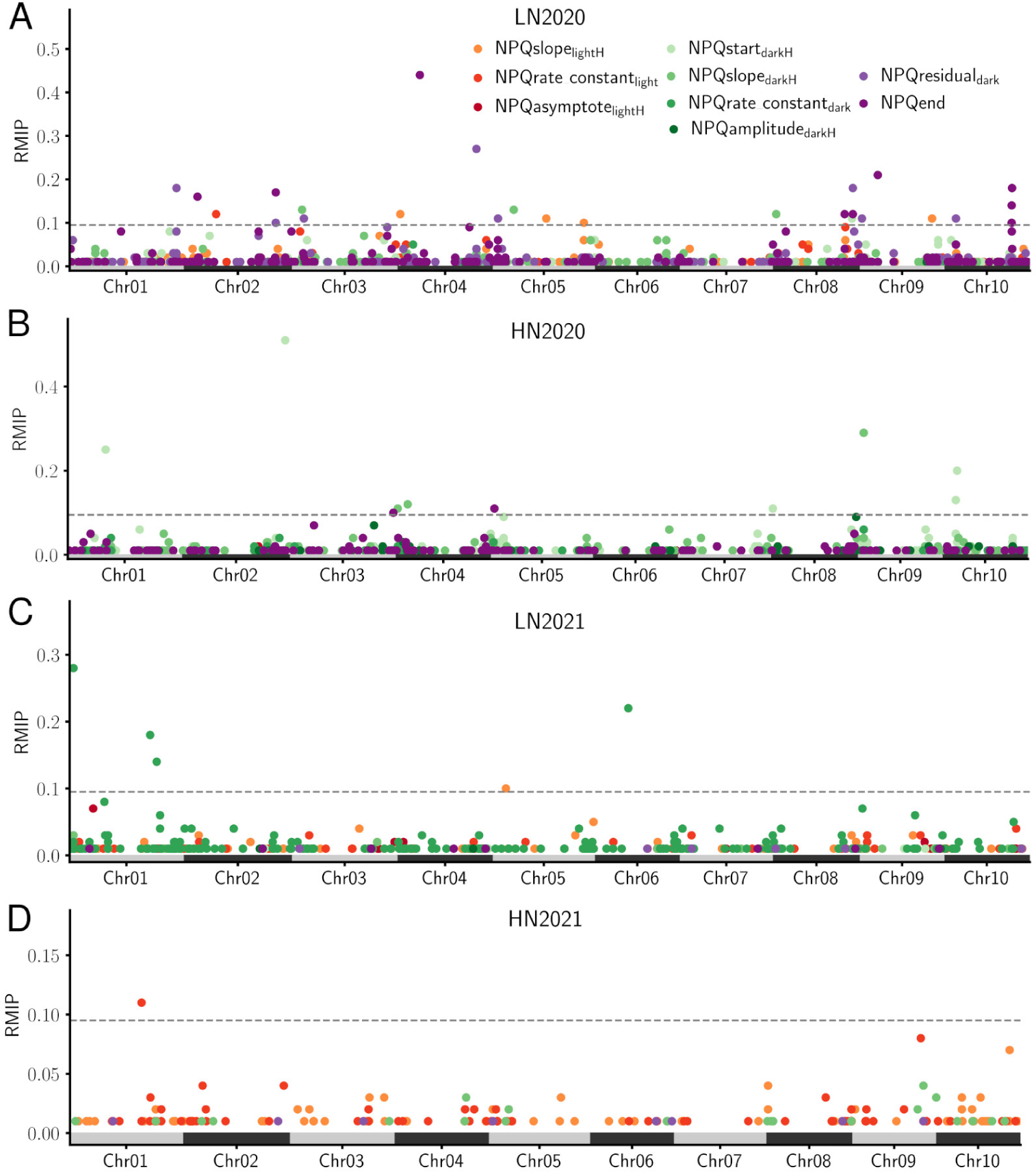
Results of GWAS for NPQ kinetic traits measured for sorghum genotypes in two treatments over two years. **A)** Results of resampling-based GWAS conducted for NPQ kinetic traits collected under low-nitrogen (LN) conditions in 2020 as shown in Figure 2A. Reproduced here for comparison with results from other treatments and years. The dashed horizontal gray line separates trait-associated markers that met the significance criteria (RMIP *≥* 0.10) from those that did not (RMIP *<* 0.10). **B)** Results of resampling-based GWAS conducted for NPQ kinetic traits collected under high-nitrogen (HN) conditions in 2020, plotted as described in panel A. **C)** Results of resampling-based GWAS conducted for NPQ kinetic traits collected under LN conditions in 2021, plotted as described in panel A. **D)** Results of resampling-based GWAS conducted for NPQ kinetic traits collected under HN conditions in 2021, plotted as described in panel A. Data used to produce this figure are given in Supplemental Data Sets S3A-D.

**Figure S5.**
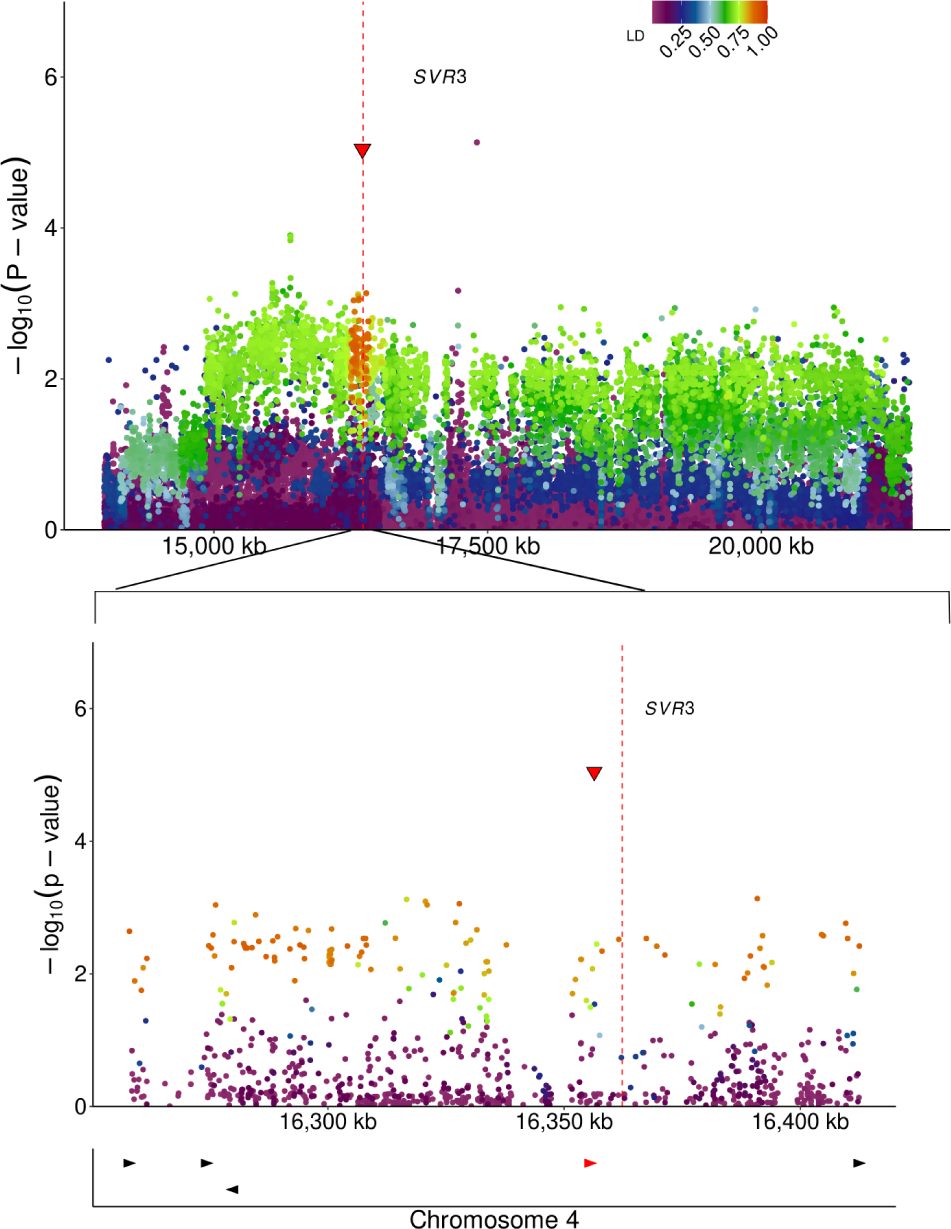
Zoomed in view of the Sobic.004G128900/*SVR3* candidate gene and LD window. Top panel, location of Sobic.004G128900 (red vertical line), the sole sorghum ortholog of *SVR3* and linkage disequilibrium (LD) of the genetic markers with Chr04:16,356,388 (red triangle)—the genetic marker identified via resampling-based GWAS for NPQend (RMIP = 0.44)—among SNPs in the interval from 14.0–21.3 Mb on sorghum chromosome 4. The color of individual points indicates the degree of LD observed between a given genetic marker and Chr04:16,356,388 among 339 re-sequenced sorghum lines. The y-axis indicates the *P*-values assigned to each marker in a mixed-linear-model-based GWAS for the *NPQ_end_*. Middle panel, zoomed-in view of the information from the top panel for the region from 16.2–16.4 Mb on sorghum chromosome 4, the interval encompassing all SNPs in LD >0.9 with Chr04:16,356,388 (red triangle). Bottom panel, the positions of all annotated sorghum gene models (red and black triangles) within the genomic interval shown in the middle panel. The red triangle indicates the position of Sobic.004G128900/*SVR3*.

**Figure S6.**
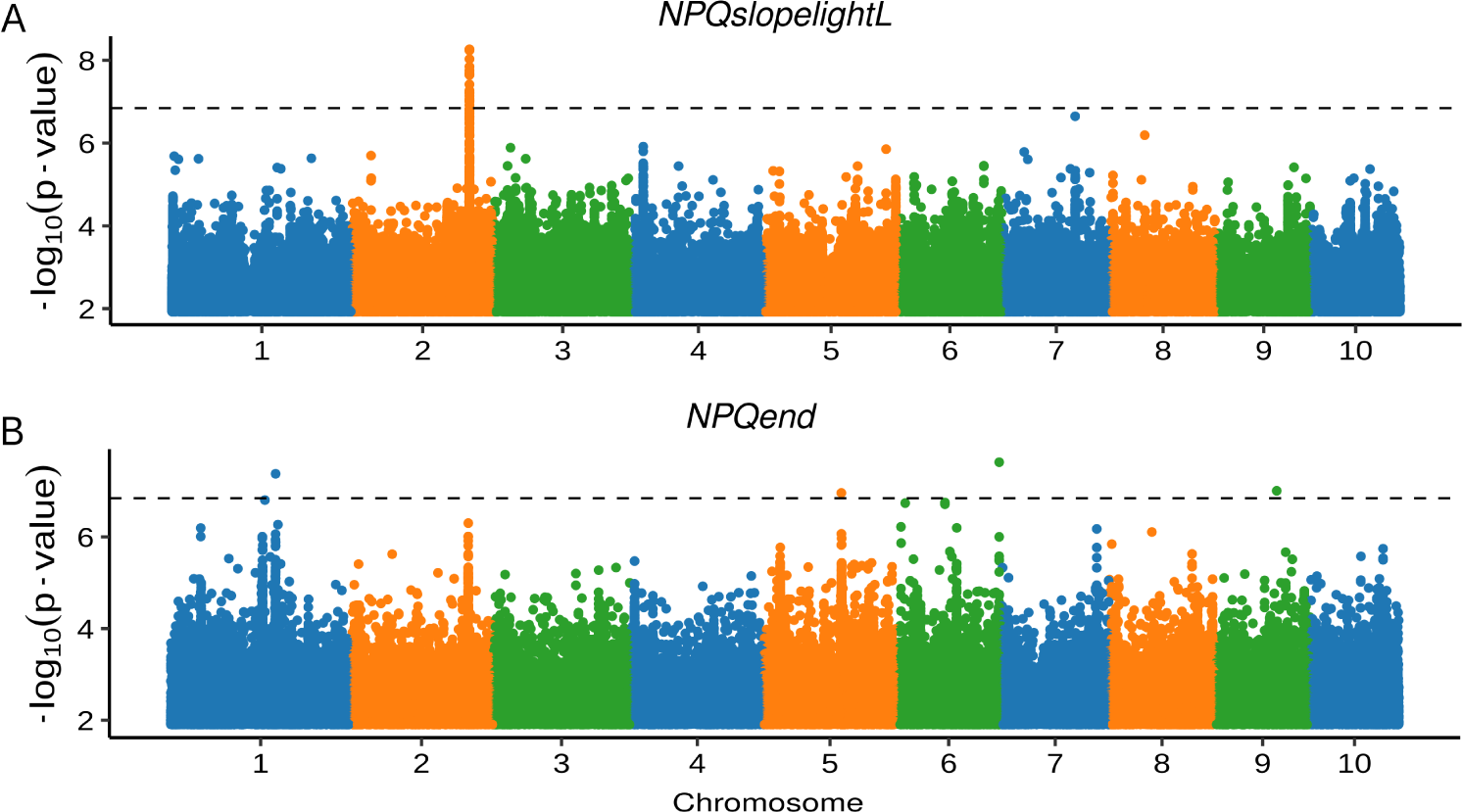
Conventional MLM-based GWAS identifies signals at the genomic interval corresponding to Zm00001eb10348 (*NYE1*). Results of a unified Mixed-Linear-Model (MLM) based GWAS implemented in rMVP (Yu *et al*. 2006; Yin *et al*. 2021). **A)** GWAS conducted for the speed of NPQ induction (*NPQslope_lightL_*) in maize in 2020 identifies a strong signal (e.g. one that exceeds the Bonferroni corrected p-value cut off) on chromosome 2 corresponding to *NYE1*. **B)** GWAS conducted for the *NPQ_end_*in maize in 2020 identifies a weak signal, (e.g. one that fails to reach the Bonferroni corrected p-value cut off) on chromosome 2 corresponding to *NYE1*.

**Figure S7.**
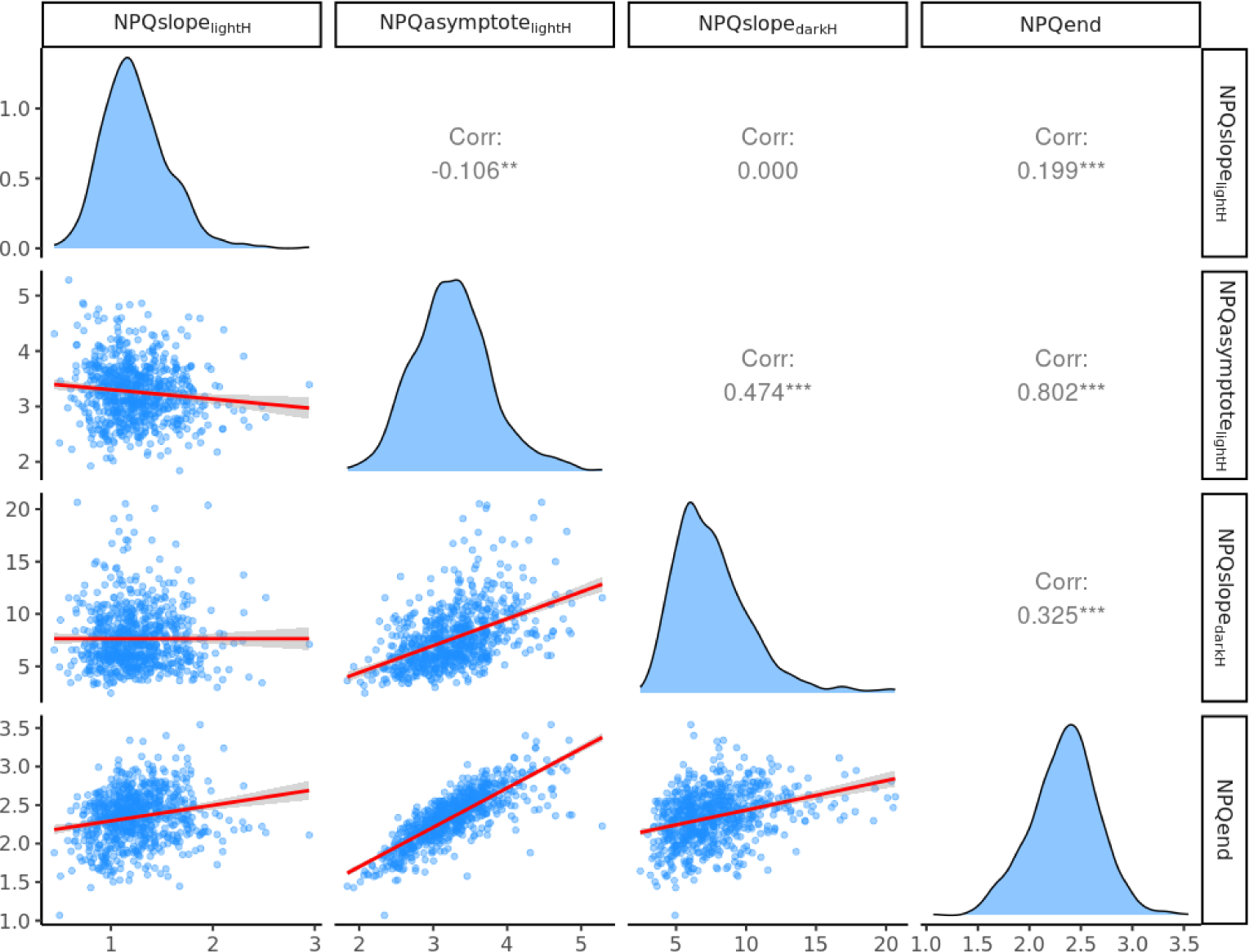
Correlation among hyperbola derived NPQ trait in maize. Pairwise correlation of published NPQ kinetics phenotypes. (*NPQslope_lighH_*, *NPQasymptote_lighH_*,*NPQslope_darkH_*, and *NPQ_end_*) from a published maize experiment conducted in 2020 (Sahay *et al*. 2023). "Corr" values presented in the top right panels are Pearson correlation coefficients between individual pairs of traits.

**Supplemental Table S1:** Locations and mapping intervals for significant GWAS hits associated with NPQ relaxation in dark traits under low nitrogen treatment in 2020.

| ID | Chr | Position | MAF | Trait(s) | RMIP | Candidate Gene | Distance | Significant in 2021 |
| --- | --- | --- | --- | --- | --- | --- | --- | --- |
| 1 | 01 | 76,062,366 | 0.46 | *NPQresidual<sub>dark</sub>* | 0.18 | N/A | N/A | No |
| 2 | 02 | 23,410,560 | 0.05 | *NPQrate*<br>*constant<sub>dark</sub>* | 0.12 | N/A | N/A | No |
| 3 | 02 | 65,900,630 | 0.47 | *NPQ<sub>end</sub>* | 0.17 | Sobic.002G274800<br>*NYE1* | 144.4 kb | No |
| 4 | 03 | 67,515,791 | 0.07 | *NPQresidual<sub>dark</sub>*<br>*NPQ<sub>end</sub>* | 0.09<br>0.07 | Sobic.003G370000<br>*PSBS* | >1 Mb | No |
| 5 | 04 | 2,395,572 | 0.36 | *NPQslope<sub>lighH</sub>* | 0.12 | N/A | N/A | No |
| 6 | 04 | 16,356,388 | 0.23 | *NPQ<sub>end</sub>* | 0.44 | Sobic.004G128900<br>*SVR3* | 1.4 kb | Yes |
| 7 | 04 | 56,606,003 | 0.18 | *NPQresidual<sub>dark</sub>* | 0.27 | Sobic.004G210500<br>*OEP37* | 555.8 kb | Yes |
| 8 | 08 | 57,327,356 | 0.09 | *NPQresidual<sub>dark</sub>*<br>*NPQ<sub>end</sub>* | 0.18<br>0.12 | N/A | N/A | No |
| 9 | 09 | 12,388,755 | 0.05 | *NPQ<sub>end</sub>* | 0.21 | N/A | N/A | No |
| 10 | 10 | 48,559,733 | 0.16 | *NPQ<sub>end</sub>* | 0.18 | N/A | N/A | No |

**Supplemental Table S2:**
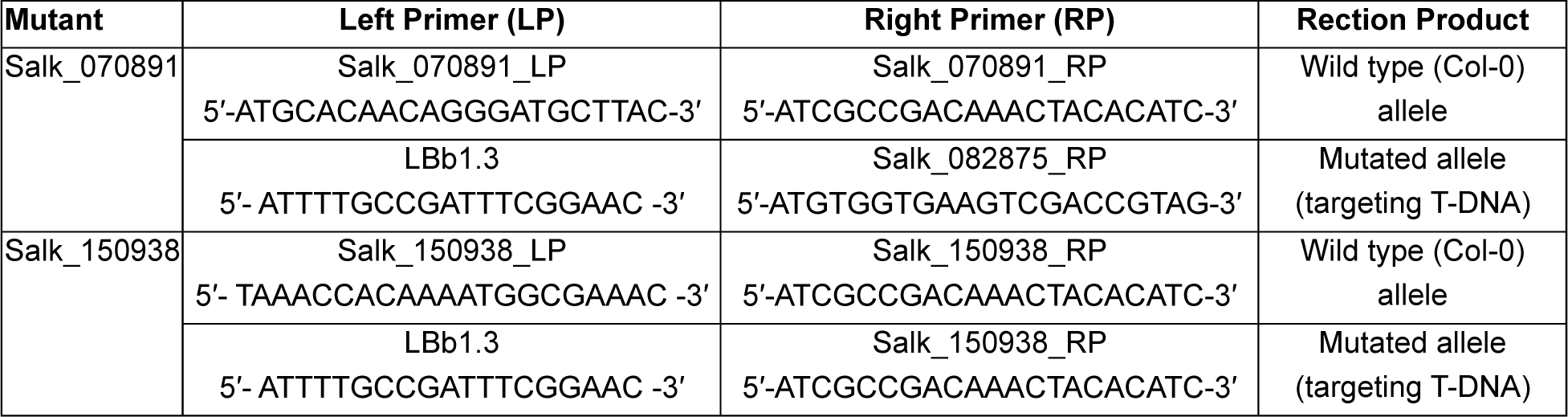
Names and sequences of primers used for PCR to verify homozygosity of Arabidopsis mutants in *NYE1* (At4g22920) gene. The Salk_070891 and Salk_150938 are the T-DNA insertional mutants in the Col-0 background.

